# Polyadenylation landscape of *in vivo* long-term potentiation in the rat brain

**DOI:** 10.1101/2024.06.05.597605

**Authors:** Natalia Gumińska, Francois P. Pauzin, Bożena Kuźniewska, Jacek Miłek, Patrycja Wardaszka, Paweł S. Krawczyk, Seweryn Mroczek, Sebastian Jeleń, Patrick U. Pagenhart, Clive R. Bramham, Andrzej Dziembowski, Magdalena Dziembowska

## Abstract

Local protein synthesis in neurons is vital for synaptic maintenance and plasticity, yet the regulatory mechanisms, particularly cytoplasmic polyadenylation, are not fully understood. This study employed nanopore sequencing to examine transcriptomic responses in rat hippocampi during *in vivo* long-term potentiation (LTP) and in synaptoneurosomes after *in vitro* stimulation. Our long-read transcriptomic dataset allows for detailed analysis of mRNA 3′-ends, poly(A) tail lengths, and nucleotide composition. We observed dynamic shifts in polyadenylation site preference post-LTP induction, with significant poly(A) tail lengthening restricted to transcriptionally induced mRNAs. The poly(A) tails of these genes showed increased non-adenosine abundance. In synaptoneurosomes, chemical stimulation led to shortening of poly(A) tails on preexisting mRNAs, indicating translation-induced deadenylation. Additionally, we discovered a group of neuronal transcripts with poly(A) tails abundant in non-adenosine residues. These tails are semi-templated and derived from extremely adenosine-rich 3′UTRs. This study provides a comprehensive overview of mRNA 3′-end dynamics during LTP, offering insights into post-transcriptional regulation following synaptic activation of plasticity in neurons.

## Introduction

Synaptic plasticity, the activity-dependent modification of synaptic efficacy, is critical for learning and memory processes^1^. In mammalian brain, long-term potentiation (LTP) and long-term depression (LTD) are canonical models for strengthening and weakening of synaptic transmission, respectively. LTP of excitatory synapses in the hippocampus is divided into early and late phases, distinguished by their duration and underlying mechanisms. After NMDA receptor-dependent LTP induction, expression of early phase LTP depends on post-translational modification and trafficking of pre-existing proteins, such as AMPA receptors and does not require massive transcriptome remodeling with only selected immediate early genes being induced. Late-phase LTP, on the other hand, involves complex transcriptomic responses and *de novo* protein synthesis, which enables stable structural and functional remodeling of synapses.

Pre-mRNA polyadenylation is a 3′-end processing mechanism for the export of RNAs from the nucleus into the cytoplasm. It contributes to mRNA stability and thus acts as a master regulator of gene expression in eukaryotes^2^. The polyadenosine tail, synthesized by the canonical poly(A) polymerase at the poly(A) site (PAS), is defined by upstream and downstream *cis*-regulatory elements recognized by the cleavage and polyadenylation specificity factor^3–5^. Most mammalian genes contain multiple PASs, leading to multiple mRNA isoforms^6^. In neurons, local translation of mRNAs in dendrites and axons enables the rapid delivery of proteins involved in the regulation of synaptic maintenance, homeostasis and plasticity^7^. The interplay between alternative polyadenylation sites (APA) defining 3′UTRs, poly(A) tail length, translation, and mRNA decay is fundamental in gene expression regulation. Indeed, translational efficiency, as well as mRNA half-lives have been shown to vary more than 1,000-fold between individual transcripts, demonstrating the range and specificity of the regulation of mRNA stability and abundance^8^. Several years ago, it was postulated that the translation of a specific group of mRNAs is activated by cytoplasmic polyadenylation^9,10^. These transcripts contain cytoplasmic polyadenylation elements (CPE) – sequence motifs recognized by CPE-binding (CPEB) proteins. Initially CPEBs were shown to induce deadenylation, to store mRNAs in a dormant state with short poly(A) tails. Upon stimulation, poly(A) tails were shown to be extended, thereby activating translation. In addition to polyadenylation, several other post-transcriptional regulatory mechanisms can regulate local protein synthesis including microRNAs or pathways that influence the activity of translation factors^11–15^.

Despite the long-standing interest in poly(A) tails and the regulation of protein synthesis, methods for reliable, genome-wide estimation of poly(A) tail length have only recently been developed^16^. Initially, most studies relied on Illumina-based protocols, such as TAIL-seq and PAL-seq^17,18^. However, DNA polymerase is unable to properly bind to long homopolymer tracts (repetitive sequences) during PCR amplification, resulting in truncated estimates of poly(A) tail length. In addition, these methods require RNA fragmentation, as the sequencer can only read 200-300 base pairs at a time^16^.

Third-generation PacBio sequencing protocols, such as FLAM-seq and PAIso-Seq, eliminate the upper limits of poly(A) tail length detection and provide a complete overview of the entire mRNA molecule^19,20^. However, they are still affected by amplification bias from library preparation, which calls into question the reliability of the results. Oxford Nanopore Technologies (ONT) offers a solution with its direct RNA sequencing (DRS) protocol. Unlike previous methods of poly(A) tail length measurement, DRS does not require amplification, allowing for real-time sequencing of single RNA molecules and generating full-length, strand-specific reads^21,22^. This approach provides the most accurate poly(A) tail length estimations and, with our recently developed algorithm Ninetails, enables the analysis of nucleotide composition^23,24^. DRS can be complemented by ONT cDNA sequencing, providing higher throughput and requiring less input material, albeit with PCR-introduced biases similar to PacBio. Consequently, the long reads from cDNA sequencing facilitate the study of mRNA regulation, including PAS usage^25^.

In this study, we employed nanopore DRS and cDNA sequencing to examine the dynamics of mRNA poly(A) tail length, alternative polyadenylation, and nucleotide composition of poly(A) tails in response to *in vivo* LTP induction in the rat dentate gyrus and following *in vitro* stimulation of isolated synaptoneurosomes. By integrating these methods, we aimed to explore the relative contributions of transcriptional induction, alternative polyadenylation, and cytoplasmic polyadenylation in regulating mRNA 3’-end dynamics. Our data does not support cytoplasmic polyadenylation as a significant contributor to translation regulation, as we were unable to demonstrate poly(A) tail extension in synaptoneurosomes or *in vivo* poly(A) tail elongation, with the exception of transcriptionally induced genes 10 min post LTP. In contrast, preexisting mRNAs rather undergo a translation-dependent deadenylation similar to that observed in the majority of somatic cells. Concurrently, we identified a new class of mRNAs with composite poly(A) tails, partially derived from extremely adenosine-rich 3′UTRs. We also provided a valuable resource with many newly identified neuronal mRNA isoforms, many of which feature LTP-induced changes in APA site usage.

## Results

### Nanopore sequencing reveals expected changes in gene expression following LTP induction

To study the activity-induced changes in post-transcriptional regulatory programs in neurons, we used LTP in the rat dentate gyrus (DG) as an established *in vivo* model for synaptic plasticity. Short bursts of high-frequency stimulation (HFS) were applied to one hemisphere (ipsilateral), while the unstimulated hemisphere (contralateral) served as a control (Figure 1a). HFS of the medial perforant path input to DG (Figure 1b) resulted in a rapid and stable potentiation of the field excitatory postsynaptic potential (fEPSP) slope: 57+/-13% for 10 min and 55+/-7% for 60 min (Figure 1c) and a rise in population spike amplitude (Figure 1c insert). After *in vivo* LTP induction (10 and 60 min post-HFS), both ipsi- and contralateral dentate gyri were dissected for RNA extraction. The stimulation paradigm used is known to induce LTP lasting at least 10 hours without decrement in anesthetized rats^26^. Since successful LTP induction leads to the expression of immediate early genes (IEGs) such as *Arc* and *Fos*^27–32^, we only analyzed the samples with increased expression of these markers in the ipsilateral hemisphere (Supplementary Figure 1, 2a).

**Figure 1.**
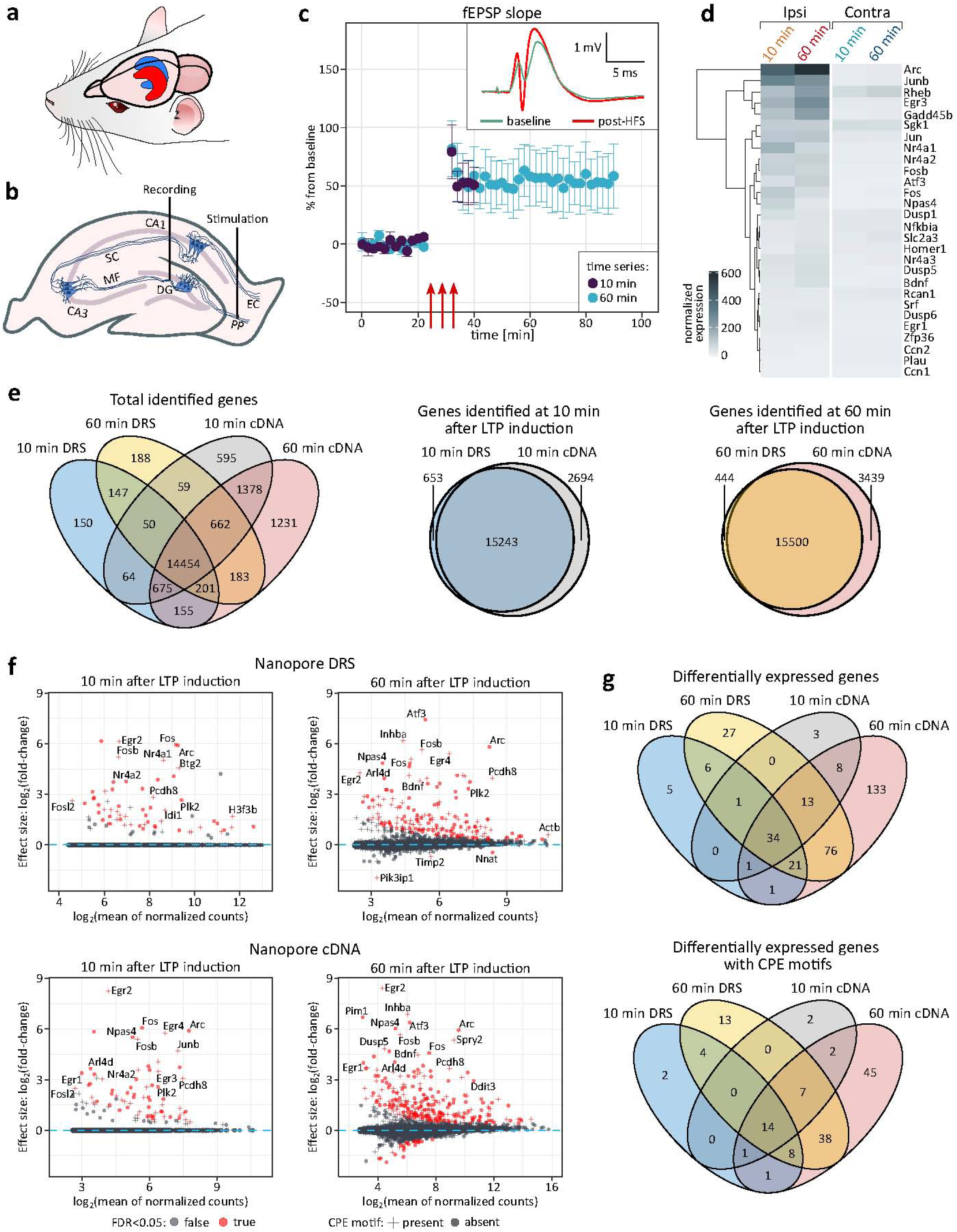
Dynamics of mRNA expression following *in vivo* hippocampal LTP induction. **a,** Schematic view of rat hippocampus. LTP was induced in the dentate gyrus of the left hippocampus (ipsilateral, red), while the right hippocampus (contralateral, blue) served as control. **b,** schematic view showing placement of electrodes for in vivo stimulation of the medial perforant path fibers from the entorhinal cortex and recording of monosynaptically, evoked fEPSPs in the dentate gyrus. DG – dentate gyrus, MF – mossy fibers, SC – Schaffer collateral, PP – perforant path, EC – entorhinal cortex. **c,** Representative HFS-induced LTP recordings in anesthetized rats. Time-course of fEPSP slope changes (10 and 60 min post-HFS) relative to baseline (meanC:±C:SEM). Red arrows indicate HFS sessions. Insert: Traces before (baseline, green) and after HFS (post-HFS, red). **d,** Expression of IEGs in the dentate gyrus following LTP (Ipsi) vs. control (Contra). Colors indicate normalized logC: expression levels. **e,** Overlap of identified genes at 10 and 60 min post-LTP using DRS and cDNA. *Left:* Venn diagram (all samples). Middle & right: Overlap at 10 and 60 min. **f,** DEGs in the dentate gyrus post-LTP. LogC: expression changes at 10 min (left) and 60 min (right) post-HFS. DEGs (red); genes with CPE motifs (+); genes without CPE motifs (dots). Results from DRS (top) and cDNA sequencing (bottom). Differential expression was calculated with DESeq2. Log_2_ fold-change estimates were shrunken (apeglm) for visualisation clarity. **g,** Overlap of DEGs at 10 and 60 min post-LTP using DRS and cDNA.

To explore the transcriptomic changes induced by neuronal activity after *in vivo* HFS, we applied two different nanopore sequencing approaches: Nanopore Direct RNA Sequencing (DRS) and Nanopore cDNA from Oxford Nanopore Technologies (ONT). Both methods produce reads encompassing whole molecules of interest, overcoming the limitations of Illumina^25^. In particular, DRS provides insights into the mRNA inventory without a synthetic proxy, avoiding amplification bias^16,22^, whereas cDNA sequencing yields higher coverage, ensuring that sufficient data is available for less expressed genes^33,34^. With DRS, we obtained an average of 0.5 million reads per sample/run, while with cDNA sequencing yielded approximately 15 million reads per sample/run (Supplementary Table 1), meeting the analysis requirements. DRS of ipsi- and contralateral dentate gyri confirmed HFS-dependent induction of IEGs, including *Arc* and *Fos,* consistent with qPCR data (Figure 1d; Supplementary Figure 1, 2a). We selected 10 and 60 min post-HFS to capture the early transcription and translation phase critical for development of late phase LTP. Te 10 min time point was chosen to captire the onset of the transcriptional responses and enable the assessment of mRNA polyadenylation of pre-existing and newly transcribed RNA. We identified 59 and 57 upregulated mRNAs at 10 min post-HFS using DRS and cDNA sequencing, respectively (Figure 1f, left panels), with no significantly downregulated transcripts. As expected, by 60 min post-HFS, gene expression changes were more pronounced, with 127 and 188 upregulated transcripts detected by DRS and cDNA sequencing, respectively (Figure 1f, right panels). The overlap between the two methods was substantial: 50% of the differentially expressed genes (DEGs) identified by DRS at 10 min were confirmed by cDNA sequencing, and at 60 min it was over 80% (Figure 1e, f, g; Supplementary Table 2, Supplementary Table 3).

Numerous specific regulatory motifs reside within mRNA sequences, predominantly in the 3′UTRs. These elements are targets for mRNA-contacting ribonucleoprotein molecules or proteins. Among them, CPEB proteins play important roles in affecting the dynamics of poly(A) tails^35,36^. Therefore, we investigated the presence of a CPE in the DEGs. Indeed, a significant fraction of upregulated transcripts harbor CPE (Figure 1f, g; see also Figure 2a, 3b).

**Figure 2.**
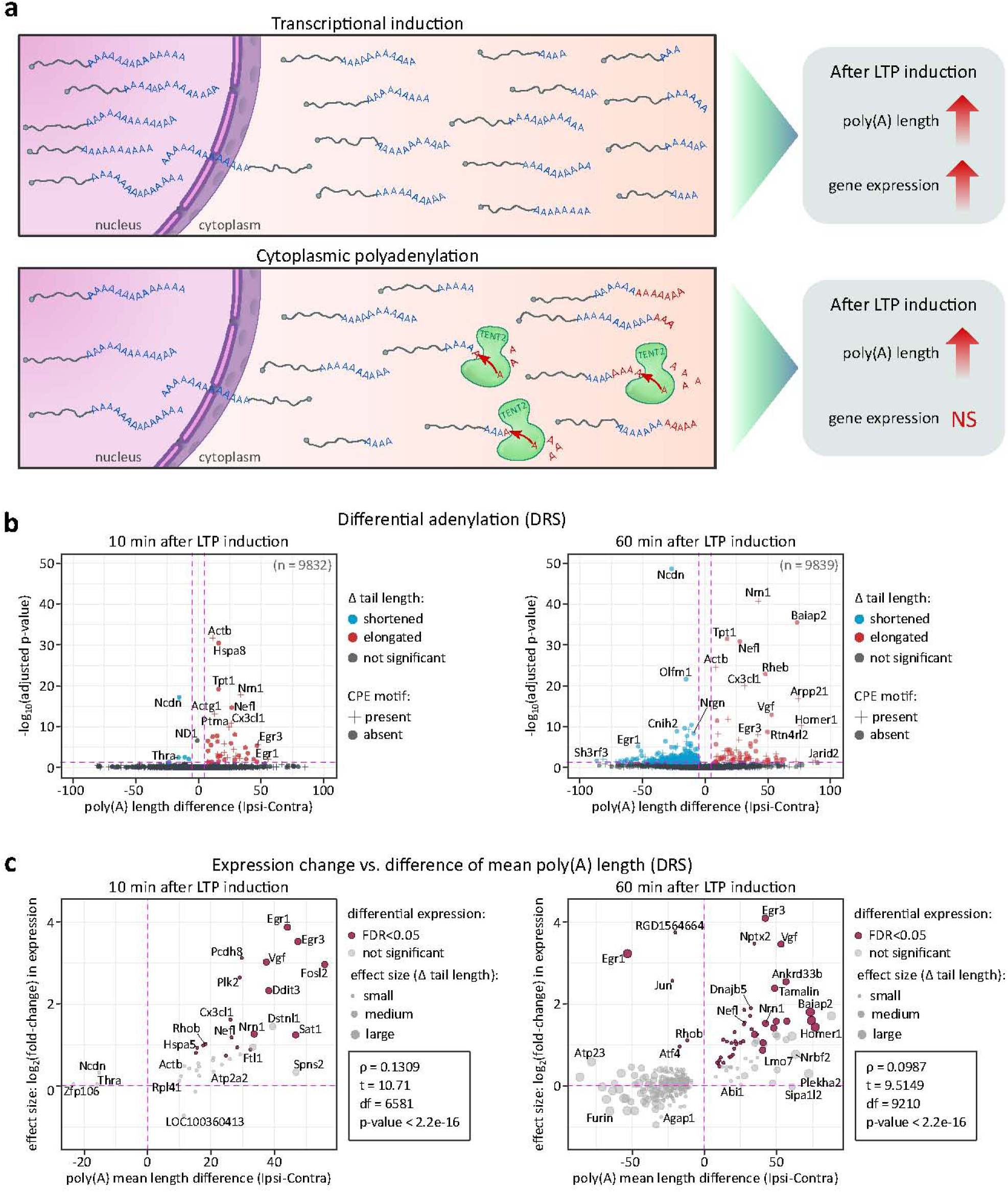
Dynamics of mRNA poly(A) tail lengths following LTP induction. **a,** Schematic of two potential mechanisms: transcriptional induction and cytoplasmic polyadenylation, underlying observed poly(A) tail length changes. **b,** Poly(A) tail length distributions at 10 and 60 min post-LTP. P.values between conditions were calculated using Wilcoxon signed-rank test, two-sided, alphaC:=C:0.05, with Benjamini-Hochberg adjustment. The pink dashed lines divide the plot area into sectors based on p-value cut-off points and differences in poly(A) tail lengths. **c,** Differential expression vs. differential polyadenylation at 10 and 60 min post-LTP. Genes with significant changes in both are highlighted (maroon). Differential expression calculated using a negative binomial generalized linear model in DESeq2. Only transcripts with statistically significant poly(A) tail length changes are displayed for clarity. Pearson correlation (two-sided) coefficient, t-test, degrees of freedom, and p.value are provided.

We conclude that LTP-inducing stimulation *in vivo* leads to the expected changes in gene expression as determined by both DRS and cDNA long-read sequencing, providing a basis for a detailed survey of the mRNA 3′-ends and poly(A) tails.

### Dynamics of mRNA poly(A) tail lengths after LTP induction

To investigate poly(A) tail dynamics in activity-dependent synaptic plasticity, we analyzed DRS and cDNA sequencing datasets, considering transcriptional induction and cytoplasmic polyadenylation as potential mechanisms for LTP-induced tail elongation (Figure 2a). While the former increases both gene expression and polyadenylation, the latter elongates tails post-transcriptionally without affecting mRNA levels.

Indeed, we observed changes in the distribution of poly(A) tail lengths, with a gradual increase in a subset of transcripts affected upon LTP (Figure 2b). Initially, 10 min post-HFS, a significant change of tail length occurred in 54 genes, of which 44 underwent tail extension. The Cohen’s d effect size of this elongation was medium for 10 genes (*Nrn1, Egr3, Sat1, Vgf, Dstnl1, Arpp21, Egr1, Fosl2, Ddit3 and Spns2*; with a mean tail length difference between ipsi-and contralateral hemispheres of 42.3 nt), small for 27 (e.g. *Actb*, *Nefl*, *Dnaja1*, *Nefm*, *Nptx1*, *Dclk1I;* with a mean tail length difference between ipsi- and contralateral hemispheres of 20 nt), and negligible for 4 genes (*Tuba1a*, *Ckb*, *Enc1*, *Hsp90aa1*; with a mean tail length difference between ipsi- and contralateral hemispheres of 7.59 nt). Among them, the most significantly enriched GO-terms were associated with synapse organization, activity, and transport (Supplementary Figure 2b). In contrast, 60 min post-HFS, a significant change in tail length occurred in 417 genes, 74 of which were elongated. The size of this effect was large for 8 genes (*Baiap2*, *Arpp21*, *Homer1*, *Nrbf2*, *Jarid2*, *Plekha2*, *Htra4*, *Mob3a*; with a mean tail length difference between ipsi-and contralateral hemispheres of 74.38 nt), medium for 22 genes (e.g. *Nrn1*, *Rheb*, *Vgf*, *Rtn4rl2*, *Egr3*, *Rbbp7*, *Synpo* and *Penk*; with a mean tail length difference between ipsi-and contralateral hemispheres of 45.64 nt), small for 40 genes (e.g. *Nefl*, *Dnaja1*, *Nefm*, *Nptx2*, *Dclk1*, *Nrp1* and *Stmn4*; with a mean tail length difference between ipsi- and contralateral hemispheres of 21.5 nt) and negligible for 4 genes (*Actb*, *Rpl41*, *Nptx1* and *Tubb4b*; with a mean tail length difference between ipsi- and contralateral hemispheres of 8.94 nt). The most enriched GO-terms were linked to cell projection and synapse organization, protein trafficking, and neurodevelopment (Supplementary Figure 2b).

We then compared changes in mRNA expression with changes in average tail length and found a strong correlation. Even at 10 min after LTP induction, poly(A) tail extension is restricted to upregulate genes arguing against cytoplasmic polyadenylation of preexisting mRNAs (Figure 2c). Many mRNAs accumulate at both time points like (e.g., *Nefl*, *Nrn1*, *Baiap2*, *Homer1*), including known IEGs (e.g., *Egf3*, *Nr4a1*) with accordingly elongated poly(A) tails. Simultaneously, a large number of mRNAs with shortened poly(A) tails at the 60 min time point correlated with their downregulation suggesting deadenylation-dependent decay. Notably, expression of *Btg2*, a general activator of deadenylation, is among the IEGs induced by the HFS (see Figure 1f)^37^. Despite the general correlation of the poly(A) tail dynamics with the differential expression, we also detected a handful of transcripts in which the elongation of poly(A) tail was independent of gene expression upregulation. Those included *Enc1* and *ActB*.

Since mRNAs bound by CPEB proteins are predicted to be the major targets of cytoplasmic poly(A) tail lengthening, we then narrowed our investigation to transcripts containing CPE (recognized by either CPEB1 and/or CPEP2,4) and compared them with the overall changes in poly(A) tail length distribution^38^ (Figure 3a, b). In general, the presence of CPE motifs did not correlate with poly(A) length distribution. Elongation was only observed on induced DEGs with the strongest effect on potential CPEB1 targets 10 min post-LTP (Figure 3b, Supplementary Figure 2c).

**Figure 3.**
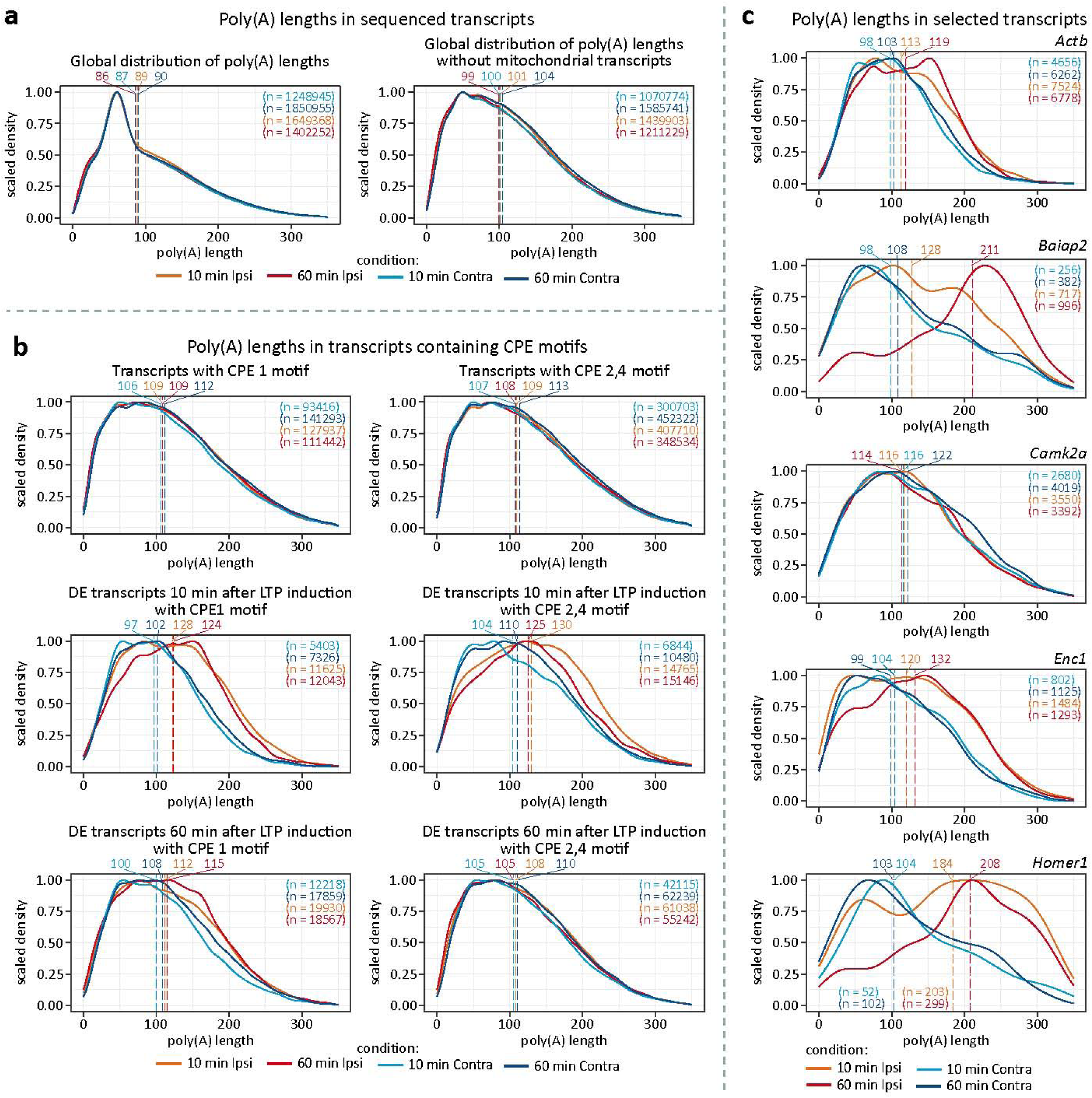
Distribution of poly(A) lengths in selected groups of mRNAs. **a**, Global distribution of poly(A) lengths: upper panel – complete datasets, bottom panel – excluding mitochondrially encoded genes. **b**, Distribution of poly(A) lengths in CPEB targets. Upper left: transcripts with CPE1 motif (total). Upper right: transcripts with CPE2,4 motifs (total). Middle left: transcripts with CPE1 motif differentially expressed 10 min after LTP induction. Middle right: transcripts with CPE1 motif differentially expressed 60 min after LTP induction. Bottom left: transcripts with CPE2,4 motifs differentially expressed 10 min after stimulation. Bottom right: transcripts with CPE2,4 motifs differentially expressed 60 min after LTP induction. All conditions are included in each panel. **c**, Distribution of poly(A) lengths in selected transcripts associated with neuroplasticity. For panels a-c, dashed lines represent median tail lengths for each condition, with median values displayed above the plotting area. The total number of reads per condition (n) is also provided.

Finally, we examined specific transcripts recognized for their significant role in hippocampal synaptic plasticity (Figure 3c), but found no evidence of cytoplasmic polyadenylation. Notably, this was also the case for *Camk2a* (Ca2^+^/calmodulin-dependent protein kinase 2-alpha), acknowledged for its activity-dependent polyadenylation (Figure 3c)^9,39^. In contrast, the transcriptionally induced *Homer1* and *Enc1* mRNAs exhibited clear tail elongation upon induction (Figure 3c). This unexpected outcome may be attributed to the relatively lower coverage provided by DRS compared to amplification-based methods. Therefore, we corroborated our findings with our cDNA data. Noteworthy, the DRS and cDNA metods are fully consistent in estimating tail length distributions (Supplementary Figure 3).

These findings demonstrate that the poly(A) tail length of certain mRNAs increases as a result of polyadenylation of the nascent mRNAs. Furthermore, none of the mRNAs with elongated tails exhibited a significant reduction in accumulation or a detectable decrease in short-tailed read abundance, which corresponds with previous studies^11^.

### Several mRNAs feature LTP-induced changes in APA site usage

Most mammalian genes contain multiple polyadenylation sites (PASs), primarily within the 3′UTR of mRNA (Figure 4a). Since 3′UTRs often harbor motifs recognized by RNA-binding proteins and miRNA target sites, changes in their length provide an efficient mechanism for regulating gene expression. Alternative polyadenylation (APA) is another post-transcriptional mechanism postulated to be pivotal in LTP^3^. Therefore, we investigated whether and how APA site usage changes following *in vivo* LTP induction in the rat brain^40,41^. To analyze APA profiles, we examined cDNA reads. Given the considerably greater sequencing depth of cDNA sequencing compared to DRS, more precise screening could be achieved. First, to eliminate nanopore-specific artifacts (e.g. fused reads), datasets were curated using pychopper. We then applied TAPAS, a software for detecting APA events in sequencing data^42^.

**Figure 4.**
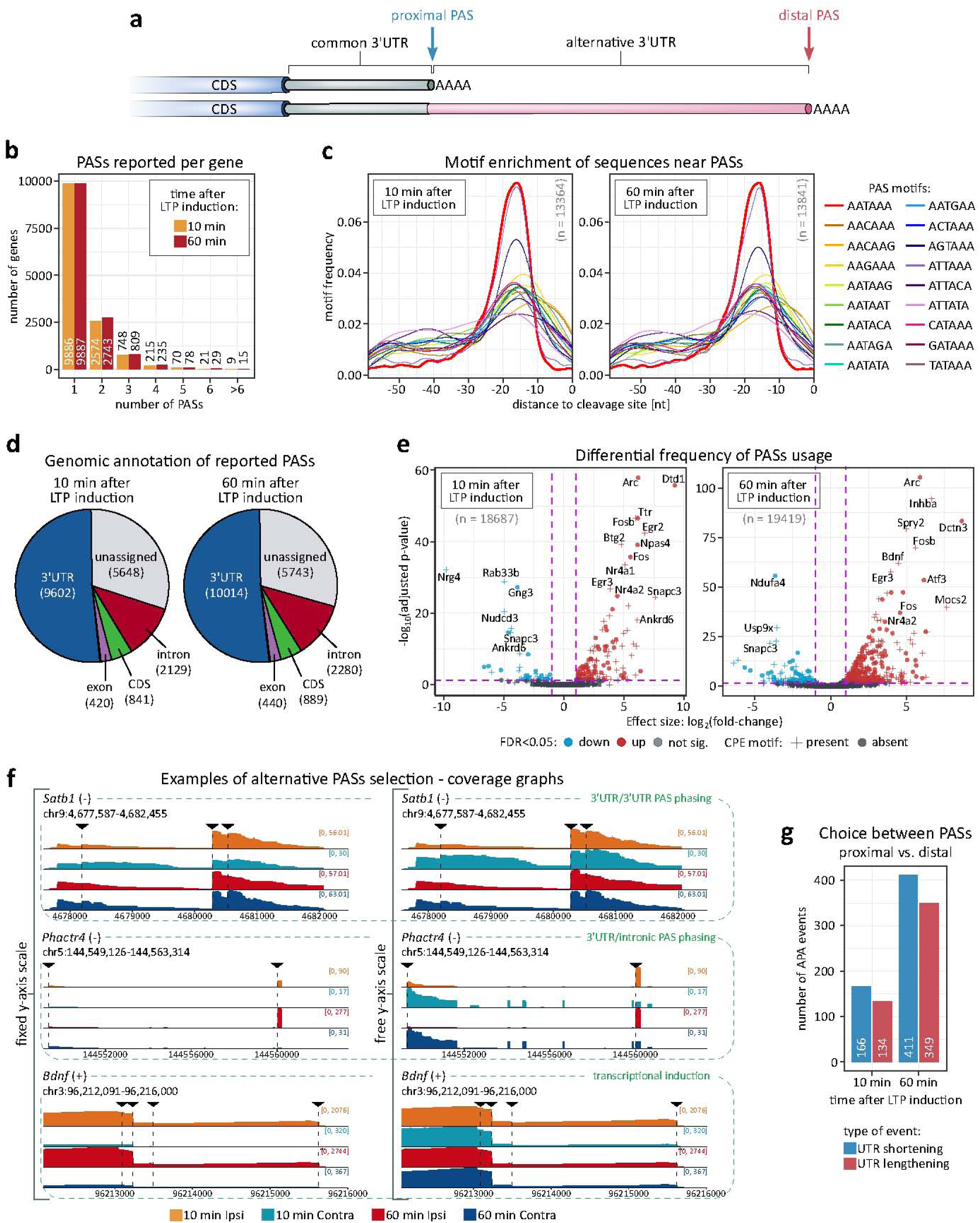
Differential polyadenylation site usage. **a**, Schematic depiction of 3′UTR containing multiple PASs. **b**, Number of PASs per gene reported by TAPAS. **c**, Enriched motifs near detected PASs. **d**, Genomic location assignment of reported PASs. **e**, Relationship between PASs usage and expression level. Expression of PASs computed with TAPAS using a negative binomial generalized linear model with the Benjamini-Hochberg adjustment (alpha= 0.1). **f**, Examples of genes exploiting multiple PASs with differential frequency due to various mechanisms (*Satb1* – alernative 3’UTRs (lengthening/shortening), *Phactr4* – intronic/3’UTR PASs selection, *Bdnf* – frequency modulated by transcriptional induction). A genome browser-like view of coverages in 3′UTR region across conditions is shown. **g**, Usage of distal and proximal PASs in genes with at least two PASs reported. A relative change in expression between conditions for proximal and distal PASs was computed in TAPAS using base-2 logarithms.

With TAPAS, we identified nearly 19,000 unique PASs in more than 13,500 genes, of which over 3,500 (∼30%) contained at least two PASs, without significant difference in the number of PASs per gene between LTP and control samples (Figure 4b). Motif enrichment analyses of sequences around PASs confirmed the significant overrepresentation of hexamer motif A[U/A]UAAA and its variants approximately 20 nt upstream of PASs (Figure 4c). Results were comparable for both control and LTP samples 10 min and 60 min post-HFS, and they were in agreement with previous studies^3,43^.

Most of the identified PASs were already known. Over 50% were located in the 3′UTR, approximately 18% within transcripts, more than 11% in introns, and ∼3% in exons. Noteworthy, nearly 30% of detected PASs were unassigned to known genomic features, meaning they appear outside annotated 3’UTRs. These likely represent novel isoforms with extended UTRs, consistent with reports demonstrating that UTRs are longer in the nervous system than in other tissues^44,45^ (Figure 4d). To refine and complement TAPAS predictions, we applied LAPA^46^, generating a high-confidence set of PASs clusters (Supplementary Figure 4). This provides a comprehensive annotation of 3′UTRs and PASs in the rat brain transcripts, including previously unreported PASs (Supplementary Figure 4a-e, Supplementary Table 4, PASs coordinates in BED format in Zenodo^47^).

Next, we analyzed 3′UTR APA events and determined whether the length of 3′UTR was changed globally following LTP induction. With TAPAS, we identified 183 differentially used PASs in 171 genes 10 min after LTP induction and 522 differentially used PASs in 452 genes 60 min post-HFS (Figure 4e). A core set of 115 genes changing expression between time points included immediate early genes, such as *Arc*, *Fos*, *Jun*, *Btg2*, *Bdnf*. GO-term analysis revealed that genes with differentially used PASs were enriched in cognition, learning and memory-related processes at 10 min post-HFS, among other terms. At 60 min post-LTP induction, they were associated with rhythmic processes, regulation of phosphorus/phosphate metabolism, and axonogenesis, along with additional enriched terms (Supplementary Figure 4g).

Multiple regulatory layers control APA, influencing mRNA stability, localization, and translation. For instance, *Satb1* undergoes 3′UTR PAS switching, with the most proximal PAS becoming less favored after stimulation, a pattern most evident after 60 min (Figure 4f, upper panel). *Phactr4* shifts toward the usage of an intronic PAS upon stimulation, promoting expression of a shorter isoform (Figure 4f, middle panel). Unlike these, *Bdnf* is an example of mRNA with multiple PASs^48^, following a transcriptional induction pattern, where the distal PAS isoform is upregulated, though the overall isoform distribution remains unchanged (Figure 4f, bottom panel).

We observed a strong link between LTP-induced changes in PAS usage and LTP-induced changes in transcript abundance (Figure 4e, Supplementary Figure 4; compare to Figure 1f). Furthermore, among all reported APA events, we noticed a general tendency to shorten the 3′UTRs in response to LTP induction, regardless of time point (Figure 4g), suggesting that proximal PASs are favored (Supplementary Table 4).

### Neuronal mRNAs with semi-templated poly(A) tails rich in non-adenosine residues

We then examined the nucleotide composition of poly(A) tails in DRS reads using our recently developed software, Ninetails^23,24^. Remarkably, in data from HFS-treated and control tissue, at both 10 and 60 min post-HFS, we observed a fraction of tails containing non-adenosines (Figure 5a, Supplementary Table 5). These transcripts commonly contained one instance of non-adenosine per tail, and only rarely two or more, coherent with our recent data (Figure 5b)^19,23^. The global proportion of decorated tails was relatively low, slightly above 3%, remaining stable across the conditions (Figure 5a, b). Subsequently, we focused on certain groups of reads: mitochondrial, DEGs 10 min post-HFS, DEGs 60 min post-HFS and potential CPEB targets (Supplementary Figure 5a). In our recent study, we found that mitochondrial transcripts have short but highly decorated tails, reflecting various fidelities of the cellular poly(A) polymerases^23,49^. Here, in data from both hemispheres, the proportion of decorated tails in mitochondrial transcripts was also relatively high and nearly equivalent to the total. It also remained stable, as did the distribution of their tail lengths (Figure 5a, Supplementary Figure 5a). Conversely, in both DEGs subsets and transcripts containing CPE motifs, we observed a higher proportion of decorated reads following LTP induction compared to the corresponding controls (Supplementary Figure 5a, b). This likely results from the activation of transcriptional mechanisms that drive the synthesis of new molecules with longer tails, where polymerase-induced errors are more frequent^23^.

**Figure 5.**
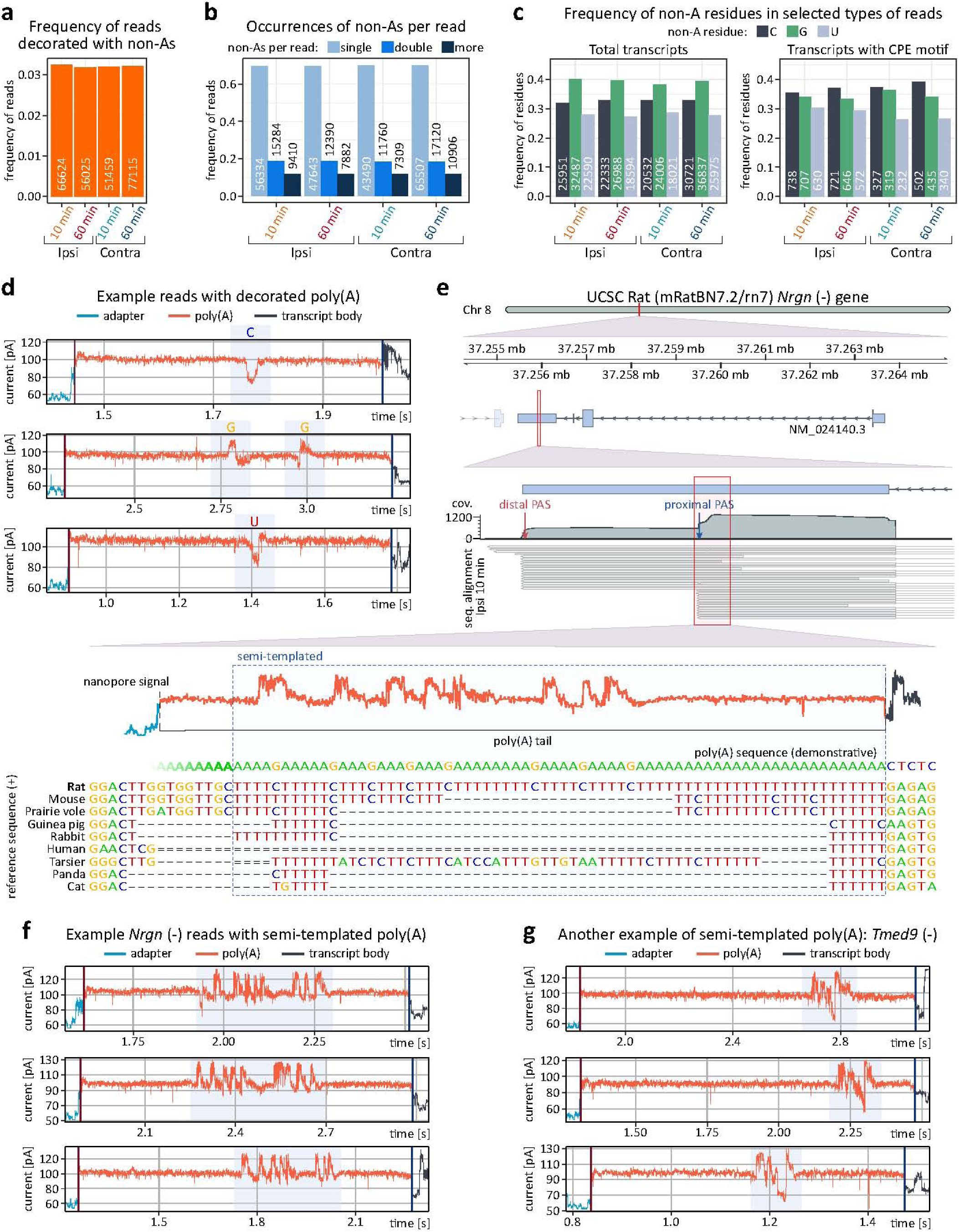
Non-adenosine residues in poly(A) tails. **a**, Total frequency of reads with poly(A) tails containing non-adenosine residues. **b**, Frequency of reads with varying numbers of non-adenosine residues per read. **c**, Frequency of non-adenosine residues (cytidine, guanosine, and uridine) in total datasets and in transcripts with CPE motifs. **d**, Example nanopore signals with decorated tails. Non-adenosine residues are highlighted. **e**, Genome browser-like plot of reads aligned to the 3’UTR of the *Nrgn* gene, with nanopore signal indicating semi-templated poly(A) tail origin in an isoform using a proximal PAS and internal non-adenosines (guanosines) in fixed positions. The reference sequence and orthologs from various organisms, as well as the reverse complement mRNA sequence, are shown. The signal is not fully aligned (resquiggled) to the sequence (it can not be done due to technical limitations of available algorithms). **f**, Nanopore signals from semi-templated tails of the *Nrgn* isoform with a shorter 3′ UTR, with fixed-position non-adenosine residues (guanosines) highlighted. **g**, Nanopore signals from semi-templated tails of the *Tmed9* gene, with fixed-position non-adenosine residues (guanosines and cytidines) highlighted. Results presented in all panels were generated using Ninetails.

We also noticed that non-adenosines are more common in longer tails (Supplementary Figure 6), indicating the possibility of their incorporation by TENT4A/B poly(A) polymerases known for their low fidelity (Supplementary Figure 5c)^49–52^. It is supported by the fact that at the global level guanosine is most prevalent, followed by cytidine and uridine (Figure 5c). This contrasts with other experimental systems, where cytidine is the most common and guanosine the least common^19,23^. Notably, gene level analysis followed by closer inspection of Ninetails output and raw nanopore signals unveiled a subset of transcripts with tails significantly enriched in non-adenosines (Figure 5d, Supplementary Figure 5d,e, Supplementary Figure 6). The most notable example is the *Nrgn* gene, where nearly 60% of the tails were decorated with (often multiple) guanosines (Figure 5e, f, Supplementary Figure 6). According to APA analysis results, *Nrgn* has two PASs (PASs coordinates in BED format in Zenodo^47^, Supplementary Table 4, Supplementary Table 5). Reads utilizing the distal PAS have blank tails (i.e. containing exclusively adenosines), whereas reads using the proximal PAS have highly decorated tails with a strikingly repetitive non-adenosine distribution pattern (Figure 5e, f). Further analysis revealed that this regularity is due to the semi-templated tail, as in this isoform the short genomically encoded 3′UTR has an extremely adenosine-rich 3′-end directly followed by poly(A) (Figure 5e, f, Supplementary Figure 6b).

In total, we identified 189 transcripts (e.g., *Bdnf*, *Camk2b*, *Homer1*, *Nrgn*, *Phactr1*, *Tmed9*) that possess semi-templated tails (Figure 5f–g, Supplementary Table 5), suggesting that the high prevalence of guanosine arises from sources other than TENT4A/B activity. Notably, the most enriched GO-terms linked to these transcripts are related to cytoskeletal reorganization, synapse structure, dendrite development, neuroplasticity, and learning (Supplementary Figure 5e). These findings suggest a previously unrecognized layer of gene regulation mediated by semi-templated tailing.

### Polyadenylation in the synaptic compartment is not induced by NMDA-R stimulation *in vitro*

Potential cytoplasmic polyadenylation in our hippocampal data might have been obscured by highly expressed transcripts from non-neuronal cells or by the predominant effect of transcriptional induction (i.e. the influx of newly synthesized long-tailed molecules; see Figure 2a)^11^. We therefore sought to identify transcripts undergoing local polyadenylation in the synaptic compartment, synaptoneurosomes stimulated chemically *in vitro* (Figure 6a; for details see Methods)^53,54^. Enrichment and stimulation of synaptoneurosomes were performed using our optimized protocol, which effectively activates translation, leading to *de novo* protein synthesis^53,54^. The effectiveness of protocol can be confirmed by western blotting (e.g. depletion/enrichment of specific markers and phosphorylation of ERK kinases; Supplementary Figure 7a, b).

**Figure 6.**
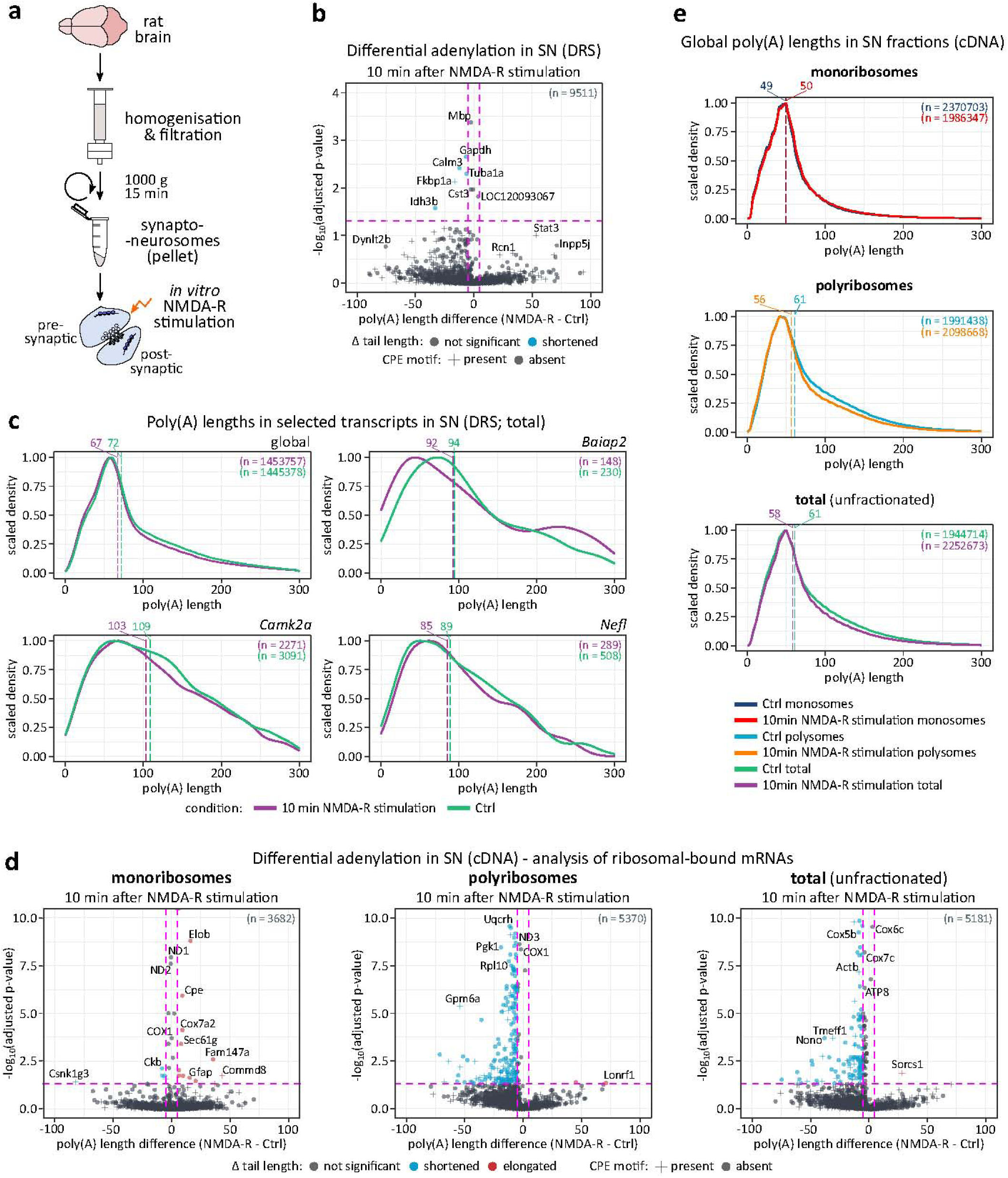
Poly(A) tail length distribution in synaptoneurosomes 10 min after *in vitro* NMDA-R stimulation. **a**, Schematic depiction of synaptoneurosomes isolation and *in vitro* stimulation. **b**, Polyadenylation profile of mRNAs in synaptoneurosomes upon NMDA-R stimulation. P.values between conditions were calculated using Wilcoxon signed-rank test, two-sided, alphaC:=C:0.05, with Benjamini-Hochberg adjustment. The dashed lines divide the plot area into sectors based on p-value cut-off points and differences in poly(A) tail lengths. Zoomed view (cut-off -log_10_(adj. p-value) = 4) highlighting statistically significant gene alterations. Most mitochondrial genes were omitted for clarity (high adj. p-value, unchanged tail length). Unscaled version of this plot is provided in Supplementary Information (Supplementary Figure). **c**, Poly(A) tail length distribution in selected groups of transcripts in synaptoneurosomes, DRS. Upper left: global (total datasets). Upper right, bottom left, and bottom right, respectively: *Baiap2*, *Camk2a,* and *Nefl* – genes related to synaptic plasticity. Dashed lines represent median tail lengths for each condition, with median values displayed above the plotting area. The total number of reads per condition (n) is also provided. **d**, Polyadenylation profile of ribosomal-bound mRNAs in synaptoneurosomes fractions upon NMDA-R stimulation. P.values between conditions were calculated using Wilcoxon signed-rank test, two-sided, alphaC:=C:0.05, with Benjamini-Hochberg adjustment. Dashed lines divide the plot area into sectors based on p-value cut-off points and differences in poly(A) tail lengths. Zoomed view (cut-off -log_10_(adj. p-value) = 10) highlighting statistically significant gene alterations. Most mitochondrial genes were omitted for clarity (high adj. p-value, unchanged tail length). Unscaled versions of these plots are provided in Supplementary Information (Supplementary Figure). **e**, Global poly(A) tail length distribution in synaptoneurosomes, fractions containing mRNA bound to monoribosomes, polyribosomes and total mRNA, cDNA sequencing. From up to down: monoribosomal, polyribosomal, and unfractionated (total) data. Dashed lines represent median tail lengths for each condition, with median values displayed above the plotting area. The total number of reads per condition (n) is also provided.

We performed DRS of mRNAs isolated from NMDA-R-stimulated synaptoneurosomes and corresponding controls to gain an accurate view of extranuclear polyadenylation (Figure 6b). As expected, we observed only mild differences in transcript abundance between the conditions (Supplementary Table 6; Supplementary Figure 7c). The poly(A) profile was also largely unchanged (Figure 6c, Supplementary Figure 7c, d). Specifically, in stimulated synaptoneurosomes no mRNAs displayed a significant shift in the poly(A) lengths from short towards longer tails that would be expected to occur if cytoplasmic polyadenylation played an important role in translational induction^8,11^. Instead, we observed a very small fraction of oligoadenylated mRNAs and a general slight shortening of poly(A) tail lengths occurring after the stimulation compared to controls. Analyzing poly(A) tail length distribution in transcripts related to synaptic plasticity also failed to confirm local cytoplasmic polyadenylation upon synaptoneurosomes stimulation (Figure 6c, Supplementary Figure 7d). There were also fewer non-adenosines in stimulated synaptoneurosomes than in corresponding controls (Supplementary Figure 7e-g). Finally, when the poly(A) tail distribution of the synaptoneurosomes and total RNA of dentate gyri were compared, the former ones exhibited considerably shorter poly(A) tails (Supplementary Figure 7h).

As a next step to better understand the relationship between poly(A) tail length and local translation in synaptoneurosomes, we performed nanopore cDNA sequencing of polyribosome-associated mRNAs. Monosomes and polysomes were separated on sucrose gradients using extracts from both NMDA-R stimulated and control synaptoneurosomes. Notably, the mRNAs associated with monosomes have significantly shorter tails than those associated with polysomes (Supplementary Figure 8a-c, Supplementary Table 6), indicating a positive relationship between poly(A) tail length and translation in synapses. However, for both monosomes and polysomes alike the total fraction, the stimulation, led to a (slight) global shortening of poly(A) tails (Supplementary Figure 8a-c), suggesting increased translation-dependent deadenylation rather than cytoplasmic polyadenylation. It further corroborated with the indication of the existence of presumably arrested polysomes and monosomes in neurons^55^.

Differential adenylation analysis comparing stimulated and unstimulated fractions of polyribosomes did not reveal any transcripts that undergo cytoplasmic polyadenylation. NMDA-R stimulation did not lead to a systematic increase in poly(A) tail lengths of any mRNAs known to be involved in synaptic plasticity in either monosome or polyribosomal fractions (Figure 6d, Supplementary Figure 8 a-c). Instead, most transcripts possessed either stable or slightly shortened poly(A) tails following stimulation (Figure 6e), indicating that local translation in synaptoneurosomes does not depend on cytoplasmic polyadenylation as a key regulatory mechanism.

## Discussion

Local protein synthesis at synapses is regulated by multiple post-transcriptional mechanisms, ensuring rapid, on-demand translation in response to neuronal activity. Given the spatial separation between the synapses and the nucleus, translational control plays a dominant role over transcriptional mRNA regulation. This is evident even in simplified *in vitro* systems, where isolated synaptic fractions exhibit robust protein synthesis activation upon stimulation^53,54^. In our study, we complemented *in vivo* LTP in the rat hippocampus with *in vitro* stimulation of synaptoneurosomes and applied long-read sequencing technologies: direct RNA and cDNA sequencing. This approach enabled an unbiased view of poly(A) tail dynamics and mRNA 3′-end processing, yielding comprehensive resources on APA site usage, tail length distribution, and its nucleotide composition.

Cytoplasmic polyadenylation has been proposed as a key mechanism controlling local translation at synapses^38^. Initially characterized in developing oocytes and early embryos^56^, this process involves the activation of dormant mRNAs with short poly(A) tails, which are elongated by cytoplasmic poly(A) polymerases, thereby rapidly initiating protein synthesis. RNA-binding proteins, particularly the CPEB family, have been implicated as major regulators of this process. However, our findings challenge this model in hippocampal neurons. We observed no systematic stimulation-induced poly(A) tail elongation in either bulk hippocampal RNA or in isolated synaptoneurosomes. Further, nanopore sequencing of ribosome-associated mRNAs revealed that NMDA-R stimulation did not induce cytoplasmic polyadenylation in translating transcripts. Instead, most poly(A) tails remained stable or exhibited slight shortening in both mono-and polyribosomal fractions. This suggests that local translation does not rely on poly(A) tail extension but is rather coupled to ongoing translation and deadenylation dynamics.

These findings also raise questions about the physiological role of noncanonical poly(A) polymerases in hippocampal neurons. While the relatively abundant TENT2 (GLD2), has been proposed to mediate CPEB-regulated cytoplasmic polyadenylation^57,58^, our data do not support its widespread activity in synaptic compartments. Similarly, TENT4A and TENT4B (GLD4), known for incorporating guanosines into poly(A) tails^49^, appear to play a minor role, as guanosine incorporation was rare except in semi-templated tails. These semi-templated transcripts, identified in our study, may represent a unique regulatory layer influencing mRNA stability and translational efficiency at synapses. Especially given that recent studies have also suggested that poly(A) tail composition, in addition to length, might play a role in gene regulation^50^. Furthermore, members of another family of cytoplasmic poly(A) polymerases, TENT5, exhibit very low expression in the hippocampus, so their impact can be ignored.

Overall, our data indicate that poly(A) tail dynamics observed in hippocampal neurons following LTP is primarily driven by transcriptional induction rather than cytoplasmic polyadenylation. The shorter poly(A) tails observed in synaptoneurosomes likely reflects the absence of recently transcribed mRNAs and active translation of quiescent, pre-existing mRNA. Instead of poly(A) tail elongation, the global activation of translation following NMDA-R stimulation correlates with mild poly(A) tail shortening, consistent with co-translational deadenylation. This is further supported by the enrichment of shorter-tailed mRNAs in synaptoneurosomes and ribosomal fractions post-stimulation. Together, these results argue against cytoplasmic polyadenylation as a major driver of local translation in hippocampal neurons. Given the growing evidence that deadenylation is a co-translational event, the global activation of translation associated with the deadenylation process explains our observation^59^.

Additionally, our analysis of alternative polyadenylation revealed widespread shifts towards proximal polyadenylation sites during *in vivo* LTP, consistent with previous reports in brain slices^3^. This preference for shorter 3′UTRs during heightened neuronal activity may enhance mRNA stability or evade miRNA-mediated repression^3,43^.

Finally, by analyzing poly(A) tail nucleotide composition, we identified a subset of mRNAs with semi-templated poly(A) tails. Many of these encode key synaptic proteins, such as neurogranin^60^. Since non-adenosine residues inhibit deadenylation, such transcripts may be particularly stable, enabling distant transport and localized translation^61^.

While further studies are needed to fully understand the regulatory impact of these modifications, our results emphasize the complexity of mRNA regulation at synapses and suggest that translation at synaptic sites is largely uncoupled from cytoplasmic polyadenylation.

## Methods

### Animals

In this study, male Sprague Dawley rats of the following age were used: (I) 5 months old (8 individuals; for the analysis of dentate gyri), 4 weeks old (26 individuals, for experiments involving synaptosome preparations). Sequenced samples/individuals are detailed in Supplementary Table 1.

The animals were housed in the laboratory animal facility under a 12 h light/dark cycle with *ad libitum* access to food and water. Experimental procedures were approved by the Norwegian National Research Ethics Committee in accordance with EU Directive 2010/63/EU, ARRIVE guidelines. All experiments were conducted by the Federation of Laboratory and Animal Science Associations (FELASA) C-certified researchers.

### Stereotaxic surgery and electrode positioning

Rats were anesthetized with urethane (1.5 mg/kg, i.p) and placed in a stereotaxic frame (David Kopfs Instruments, USA) once unresponsive to toe pinch. A thermostatically controlled heating pad was placed under the animal to maintain a temperature of 37°C. The scalp was incised longitudinally and retracted with bulldog clamps. Two burr holes were drilled rostral to bregma for ground and reference screws. For stimulation, a craniotomy was performed over the left hemisphere (7.9 mm caudal, 4.2 mm lateral, 2.5 mm ventral to bregma), targeting the angular bundle of the medial perforant path. For recording, a craniotomy was made over the same hemisphere (3.9 mm caudal, 2.3 mm lateral, ∼2.5-3 mm ventral to bregma), targeting the hilar region of the dentate gyrus. The recording electrode depth was determined when test pulses triggered the maximum field excitatory postsynaptic potential (fEPSP) slope.

### *In vivo* electrophysiology

The stimulating electrode delivered test pulses every 30 s unilaterally into the medial perforant path to evoke fEPSPs in the hilar region of the dentate gyrus, measured extracellularly using a recording electrode. Stimulus intensities (typically 100-500 µA) were set to elicit one-third of the maximum population spike. Baseline potentials were recorded for 24 min. Following baseline acquisition, the current was doubled, and high frequency stimulation (HFS) was applied to induce long-term potentiation (LTP). Simulation was delivered via a Grass S88 Dual output stimulator (Grass Medical Instruments) using a three-session paradigm (5-min intervals), totaling 10 min 30 s. Each session comprised four trains of 8 pulses at 400 Hz with 10 s inter-train intervals resulting in 96 pulses^26^. Post-HFS, the current was immediately reduced to baseline levels, and recordings continued at 0.033 Hz for either 10 min or 60 min, depending on the experimental group.

fEPSPs were recorded using Data Wave Technologies Work Bench Software (Longmont, CO). The fEPSP slope was determined as the maximum positive dV/dt within a defined time window, from onset to one-third of the positive peak, capturing early EPSPs characteristic of the monosynaptic connection from the entorhinal cortex to the dentate gyrus. Slope and population spike amplitude data were processed in Microsoft Office Excel 2010 (Microsoft Corporation, USA), averaged over 2-min bins (mean of four consecutive measurements), and normalized to the 24-min baseline mean. GraphPad Prism (San Diego, CA) was used for statistical analysis, comparing baseline to post-HFS values.

### Brain dissection

Immediately after the experiment, the rat was removed from the stereotax. For dissection: the brain was removed from the skull and placed on filter paper above a glass plate cooled with ice. Ice-cold saline solution was used periodically through dissection to cool the brain down. With the hippocampus isolated, blood vessels and connective tissue were removed before the Cornu Ammonis regions were separated from the dentate gyrus and each was put in cold microtubes. The samples were put on dry ice and kept at -80°C. This was done for both the ipsilateral and contralateral dentate gyri.

### RNA extraction and quality check

Tissue was maintained on dry ice prior to homogenization in 1 mL of Trizol using 1 mL glass/glass Dounce homogenizer. The homogenate was transferred to 1.5 ml LoBind type Eppendorf tube and centrifuged at 12,000 g for 5 min at 4°C to remove lipid content. RNA was then extracted from the supernatant following the manufacturer’s protocol. Precipitation was performed by adding glycogen (GlycoBlue Coprecipitant, 15 mg/mL; 30 µg per sample), 1 volume of isopropanol, and incubating overnight at -20°C. The RNA pellet was dissolved in Rnase-free dd H_2_O and initially assessed for quality using a Nanodrop 2000 spectrophotometer, yielding ∼20µg per sample. Further purification was performed using KAPA Pure Beads (KAPA Biosystems) according to manufacturer’s instructions. RNA integrity number (RIN) was determined using an Agilent Bioanalyzer (Agilent RNA 6000 Nano Kit). For synaptoneurosomes, RNA was isolated using the same method, with ∼1 mL Trizol per 100 mg pellet.

### qRT-PCR

1 µg of RNA samples were reverse transcribed using random primers (GeneON, Ludwigshafen am Rhein, Germany; #S300; 200 ng/RT reaction) and SuperScript IV Reverse Transcriptase (Thermo Fisher Scientific). Next, the cDNA samples were amplified using specific primers:

c-fos Fwd: 5′-CCCATCCTTACGGACTCCC-3′

c-fos Rev: 5′-GAGATAGCTGCTCTACTTTGCC-3′

Arc Fwd: 5′-AAGTGCCGAGCTGAGATGC-3′

Arc Rev: 5′-CGACCTGTGCAACCCTTTC-3′

Gapdh Fwd: 5′-ATCAAGAAGGTGGTGAAGCA-3′

Gapdh Rev: 5′-CATACCAGGAAATGAGCTTC-3′).

Each reaction was set in a final volume of 15 μl, using PowerUp SYBR Green Master Mix (#A25742 Thermo Fisher Scientific) in a LightCycler480 (Roche). Fold changes in expression were determined using the ΔΔ Ct (where Ct is the threshold cycle) relative quantification method. Values were normalized to the relative amounts of *Gapdh* mRNA.

### Nanopore direct RNA sequencing

The DRS was performed as described by Bilska et al.^62^. Briefly, for dentate gyri, 4.5-5 μg of total RNA was mixed with 250 ng of oligo(dT)-enriched RNA from *Saccharomyces cerevisiae* and 5 ng of *in vitro* transcribed poly(A) standards. For synaptoneurosomes, 3 μg of cap-enriched RNA was mixed with 200 ng of *S. cerevisiae* oligo(dT)-enriched RNA and 5 ng of poly(A) standards. Libraries were prepared using the Direct RNA Sequencing Kit SQK-RNA002 (ONT) according to the manufacturer’s instructions. Sequencing utilized FLO-MIN106D (9.4.1) flow cells on a MinION device controlled by MinKNOW software (ONT). Basecalling was performed with Guppy (ONT).

### Nanopore cDNA sequencing

For dentate gyri, 200 ng of total RNA was used to prepare nanopore cDNA sequencing libraries with the cDNA-PCR Sequencing Kit SQK-PCS111 (ONT) according to the manufacturer’s instructions. Sequencing was carried out using FLO-MIN106D (9.4.1) flow cells on a MinION device controlled by MinKNOW software (ONT). The basecalling was performed with Guppy (ONT) and Dorado (ONT).

For synaptoneurosomes fractionated with linear sucrose gradient, 500 ng of RNA isolates was used to prepare libraries with PCR-cDNA Barcoding Kit SQK-PCB111-24 (ONT) according to the manufacturer’s instructions. Sequencing was performed using FLO-MIN106D (9.4.1) flow cell with the MinION device controlled by MinKNOW software. Obtained reads were basecalled and poly(A)-profiled using Dorado (ONT).

### Poly(A) tail length profiling

Basecalled RNA reads were mapped to mRatBN7.2-derived reference transcriptome using minimap2^63^ (-k 14 -ax map-ont --secondary = no) and processed using samtools^64^ to filter out supplementary alignments and reads mapping to the reverse strand (samtools view -b -F 2320). The lengths of the poly(A) tails were estimated using the Nanopolish polya function^22^. Only reads tagged by Nanopolish as PASS or SUFFCLIP were considered in the following analyses.

The lengths of poly(A) tails in cDNA reads were estimated during basecalling with dorado (--estimate-poly-a), extracted from the source files, and aggregated in tabular form.

The NanoTail package^65^ was used to perform statistical analysis. The p-values were calculated using Wilcoxon signed-rank test (two-sided, alpha = 0.05) for transcripts represented by 10 or more reads, and adjusted for multiple comparisons with Benjamini-Hochberg method (alpha = 0.05). Cohen d was used as an effect size measure. For values x<0.2, effect size was considered ’negligible’, for values 0.2≤x<0.5 it was considered ’small’, for values 0.5≤x<0.8 it was considered ’medium’, and for values x≥0.8, it was considered ’large’. Transcripts were considered significantly altered in poly(A) tail length if the adjusted p-value was less than 0.05 and the difference in tail length was greater than 5 nt between the hemispheres.

### Differential expression

Reads were mapped to the mRatBN7.2 reference genome using minimap2^63^ (-ax splice -- secondary = no -uf). The read distribution across genes was summarized using featureCounts from the Subread package^66^ (-L --fracOverlap 0.5 --fracOverlapFeature 0.5 -s 1). Reads overlapping multiple features and multimappers were excluded. Differential expression analysis was performed using DESeq2^67^ with default settings. For visualisation clarity, log_2_ fold-change estimates were shrunken using apeglm^68^ method.

### Nucleotide composition of poly(A) tails

Non-adenosine nucleotides in DRS reads were assessed with Ninetails^23^ (check_tails function; pass_only = FALSE, qc = TRUE). The raw classification results were further processed and visualized using Ninetails postprocessing functions with default parameters.

The last 60 nucleotides of each annotated 3’UTR were extracted from the mRatBN7.2 FASTA reference based on the GCF_015227675.2 reference annotation. Antisense strand sequences were reverse-complemented, and A/T percentages were calculated. Transcripts meeting at least one of the following criteria were retained: (I) A/T content >60% (147 transcripts) or (II) ≥13 consecutive A/T nucleotides, with the last 3 also being A/T (406 transcripts). The 13-nucleotide threshold was based on the R9 pore pentamer reading context, covering 2.5 independent nucleotide contexts – sufficient for detecting meaningful signal differences. After merging and deduplication, 471 transcripts with potential semi-templated tails were identified.

A 60-nucleotide upstream context was extracted for PAS sites predicted by LAPA and TAPAS. Applying the same filtering criteria resulted in 518 transcripts after deduplication. After merging reference-based (471) and PAS-predicted (518) sets, 988 transcripts remained. Of these, only those with ≥10 DRS reads and ≥3 poly-A tails with non-adenosine nucleotides were retained, yielding 189 transcripts potentially possessing semi-templated tails.

Analysis was performed in R using GenomicRanges^69^, IRanges^69^, Biostrings^70^, stringr^71^, dplyr^72^, and Ninetails^23^.

### Alternative polyadenylation

The cDNA reads were preprocessed using pychopper (ONT). Reads fulfilling quality criteria and rescued fused reads were combined, then mapped to the mRatBN7.2 reference genome with minimap2^63^ (-ax splice --secondary = no -uf -k 14). Alternative polyadenylation site analysis was performed with TAPAS^42^ and LAPA^46^.

For the analysis using the TAPAS software, the mRatBN7.2 NCBI reference annotation (GCF_015227675.2-RS_2023_06) [https://www.ncbi.nlm.nih.gov/datasets/genome/GCF_015227675.2/] was converted into the refFlat format using the gtfToGenePred script from UCSC tools [http://hgdownload.soe.ucsc.edu/admin/exe/linux.x86_64/]. Since TAPAS requires all gene identifiers (column 2) to begin with the “NM_” prefix [https://github.com/arefeen/TAPAS/issues/2], this modification was applied manually in R environment. Additionally, the columns were reordered according to the format specified by the software developers.

For the analysis using the LAPA software, the same NCBI reference was refined using scripts from AGAT^73^. First, intron coordinates were incorporated into the annotation using the agat_sp_add_introns.pl script. The resulting file was then converted to GTF format using the agat_convert_sp_gff2gtf.pl script. Since the LAPA package requires UTR sequences to be labeled in a non-standard format (“five_prime_utr” and “three_prime_utr”), the annotation was further modified in terminal (sed -e ’s/<5UTR\>/five_prime_utr/g’ -e ’s/<3UTR\>/three_prime_utr/g’ [input file name] > [output file name], where the placeholders in square brackets should be replaced with the actual file names). All resulting custom files are provided on Zenodo^47^. The raw predictions of LAPA were further processed in Python according to the instructions provided by LAPA developers [https://colab.research.google.com/drive/1QzMxCRjCk3i5_MuHzjozSRWMaJgdEdSI?usp=sharing].

Differential usage of poly(A) sites was calculated using Fisher’s exact test (two-sided, alpha = 0.05). P-values were adjusted using the Benjamini-Hochberg method. Given PAS was considered as differentially expressed if adjusted p-value was less or equal to 0.1.

### CPE motif search

blished CPE1 and CPE2,4 motif sequences represented as position-dependent letter-probability matrices^74^ were used to identify CPE motifs, recognized by CPEB1 and CPEB2-4, respectively. Motif searches were conducted separately using the FIMO software from the MEME suite^75^ (parameters: --bgfile --nrdb thresh 1.0E-4).

### Synaptoneurosomes isolation and *in vitro* stimulation

Synaptoneurosomes were prepared as described previously^54^. Briefly, Krebs buffer (2.5 mM CaCl_2_, 1.18 mM KH_2_PO_4_, 118.5 mM NaCl, 24.9 mM NaHCO_3_, 1.18 mM MgSO_4_, 3.8 mM MgCl_2_, and 212.7 mM glucose) was aerated at 4°C for 30 min using an aquarium pump, then adjusted to pH 7.4 with dry ice. The buffer was supplemented with 1× protease inhibitor cocktail (EDTAC:free, Roche) and RNase Inhibitor (RiboLock, 60 U/ml, Thermo Fisher Scientific).

Rats were euthanized by cervical dislocation, and hippocampi were dissected. Tissue from one hemisphere was homogenized in a 1.5 ml of Krebs buffer using a Dounce homogenizer (10–12 strokes). To prevent the stimulation of synaptoneurosomes, all steps were performed on ice. Homogenates were passed gravitationally through presoaked nylon mesh filters (100, 60, 30, and 10 μm, Merck Millipore) into a 50C:ml polypropylene tube, centrifuged at 1,000 g for 15 min at 4°C, washed, and resuspended in Krebs buffer with protease and RNase inhibitors.

*In vitro* stimulation of NMDA receptors on synaptoneurosomes was performed as described before^54^. Briefly, freshly isolated synaptoneurosomes were prewarmed at 37°C for 5 min, stimulated with 50 μM NMDA and 10 μM glutamate for 30 s, followed by 120 μM APV and incubation at 37°C for 10 min. Unstimulated samples kept on ice served as controls. Synaptoneurosomes from 5 rats were pooled and split into control and stimulated samples to produce enough material for Nanopore Direct RNA Sequencing (DRS). Four independent isolations were performed for DRS.

### Polysome profiling

Linear sucrose gradients (10–50%) were prepared in GB buffer (10 mM HEPES–KOH pH 7.2, 150 mM KCl, 5 mM MgCl2, Protease Inhibitor Cocktail (Roche), 100 μg/ml cycloheximide (CHX; Thermo Fisher Scientific), 4 U/ml RiboLock (Thermo Fisher Scientific), and nuclease-free water). Krebs buffer was prepared as described in ‘Synaptoneurosomes isolation and *in vitro* stimulation’ section.

Rats were sacrificed by cervical dislocation. Brains were rapidly harvested, briefly chilled in ice-cold Krebs buffer, and hippocampi were dissected and halved transversely. Tissue from one half was homogenized in 1.5 ml Krebs buffer using glas-glas Dounce homogenizer (10-12 strokes), and homogenates were pooled. All steps were performed on ice to prevent synaptoneurosome stimulation. Homogenates were further processed as described in the ‘Synaptoneurosomes isolation and *in vitro* stimulation’ section, and resuspended in 1 ml of Krebs buffer with protease and RNase inhibitors.

Cycloheximide (100µg/ml) was added to control synaptoneurosomes kept on ice. The second aliquot was prewarmed at 37°C (800 rpm, thermoblock) for 5 min, stimulated with 50 μM NMDA and 10 μM glutamate for 30 s, followed by 120 μM of APV, and incubation for 10 min at 37°C (800 rpm, thermoblock). Stimulation was halted by adding cycloheximide (100µg/ml) and incubating on ice for 2 min. Samples were centrifuged at 1,000 g for 5 min at 4°C, and synaptoneurosomes were resuspended in GB buffer with 1,5% NP-40. After 10 min on ice, samples were centrifuged at 20,000 g for 15 min. RNA was extracted by mixing 300 μl of lysate with 900 μl of Tri reagent LS.

Equal volumes of samples (1,1 ml) were loaded onto sucrose gradients, and ultracentrifuged at 38,000 rpm for 2 h at 4°C using an Optima XPN ultracentrifuge with an SW41Ti rotor (Beckman Coulter). Sucrose fractions were collected using a Density Gradient Fractionation System (Teledyne ISCO) with a Foxy Jr. Fraction Collector. Fractions were stored at -80°C for further analysis. Absorbance at 254 nm was recorded during collection to generate RNA distribution profiles.

### Western blotting analysis of synaptoneurosomal preparations

Equal amounts of protein from homogenate and synaptoneurosomal fraction were resolved on SDS-PAGE (10%, TGX Stain-Free FastCast Acrylamide Solutions, BioRad). After electrophoresis proteins in the gel were visualized using Bio-Rad’s ImageLab software to verify the equal protein loading. Proteins were transferred to PVDF membranes (pore size 0.45 µm, Immobilon-P, Merck Millipore) using the Trans-Blot Turbo Blotting System (BioRad). Membranes were blocked for 1 h at room temperature in 5% non-fat dry milk in PBS-T (PBS with 0.01% Tween-20), followed by overnight incubation at 4°C with primary antibodies (catalog numbers and producers of antibodies are listed in Supplementary Table 1) in 5% milk in PBS-T. Blots were washed 3 × 5 min with PBS-T, incubated 1 h at room temperature with HRP-conjugated secondary antibody (1:10 000 in 5% milk) and washed 3 × 5 min with PBS-T. In the case of MAPK/Phospho-MAPK Family Antibody Sampler Kit, membranes were blocked for 1 h at room temperature in 5% bovine serum albumin (BSA) in PBS-T, primary and secondary antibodies were diluted also in 5% BSA in PBS-T. HRP signal was detected using Amersham ECL Prime Western Blotting Detection Reagent (GE Healthcare) on ChemiDoc Imaging System (BioRad) using automatic detection settings.

### Statistics and reproducibility

Sample size was not pre-determined using a statistical method. Statistical analyses were conducted on data from two or more biologically independent replicates. Quantitative data were analyzed within the R environment, with specific statistical tests detailed in the figure legends. Normality was assessed using the Shapiro-Wilk test. Most experiments were performed at least twice, producing consistent results.

### Data visualization

The visualization of the data was performed in the R environment. Poly(A) nucleotide composition analyses using Ninetails^23,24^. Heatmaps were created with ComplexHeatmap^76^, and genome browser-like diagrams were drawn using ggcoverage^77^. Gene ontology enrichment was performed using g:Profiler^78^ and ClusterProfiler^79^. The remaining diagrams were created using ggplot2^80^.

## Data availability

Raw sequencing data (fast5 files) are deposited at the European Nucleotide Archive (ENA) [https://www.ebi.ac.uk/ena/browser/home] under project accession numbers PRJEB75352 and PRJEB75356 with sample numbers listed in Supplementary Table 1. Source data used for plotting (including statistics) are provided in Supplementary Tables 2-6.

Supplementary Tables and other resources (e.g. custom references, CPE motifs, APA site coordinates) are deposited in Zenodo^47^ [https://doi.org/10.5281/zenodo.11478468].

Supplementary Figures are provided in the Supplementary Information file. Additional files/code snippets and information are available from the authors upon request.

## Code availability

Custom software (NanoTail) used for poly(A) tail analysis in R is available on Github repository [https://github.com/LRB-IIMCB/nanotail].

Custom software (Ninetails) used for non-adenosine profiling in R is available on Github repository [https://github.com/LRB-IIMCB/ninetails] and in Zenodo repository [https://doi.org/10.5281/zenodo.11478467].

Other programs, tools and scripts used in this work are listed in Supplementary Table 1.

## Acknowledgements

We would like to thank all the lab members of our collaborating groups for their support and fruitful discussions. The research leading to these results was funded by the Norwegian Financial Mechanism 2014 -2021, no. UMO-2019/34/H/NZ3/00733.

## Author contributions statement

NG supported by PK performed all the bioinformatic analyses and prepared figure drafts; FPP supported by PUG performed *in vivo* LTP. BK, PW and JM analyzed RNA samples and performed experiments on synaptoneurosomes. SM and SJ were responsible for nanopore sequencing. NG, FPP, AD, and MD wrote the manuscript. CRB, AD, and MD jointly supervised and directed the studies.

## Description of provided supplementary data

**Supplementary Table 1. Key resource table. (A)** Characteristics of analyzed material (e.g. sample identifiers, number of animals used, reads produced, accession numbers). **(B)** Key resources (antibodies, reagents, software).

**Supplementary Table 2. Summary of dentate gyri DRS data per gene. (A)** Differential expression and differential adenylation data for 10 min timepoint. **(B)** Differential expression and differential adenylation data for 60 min timepoint. **(C)** GO-terms for genes with significantly elongated poly(A) tails in 10 min timepoint. **(D)** GO-terms for genes with significantly elongated poly(A) tails in 60 min timepoint. **(E)** GO-terms for upregulated genes in 10 min timepoint. **(F)** GO-terms for upregulated genes in 60 min timepoint. **(G)** GO-terms for upregulated genes with CPEB-binding motifs in 10 min timepoint. **(H)** GO-terms for upregulated genes with CPEB-binding motifs in 60 min timepoint.

**Supplementary Table 3. Summary of dentate gyri cDNA data per gene. (A)** Differential expression and differential adenylation data for 10 min timepoint. **(B)** Differential expression and differential adenylation data for 60 min timepoint. **(C)** GO-terms for genes with significantly elongated poly(A) tails in 10 min timepoint. **(D)** GO-terms for genes with significantly elongated poly(A) tails in 60 min timepoint. **(E)** GO-terms for upregulated genes in 10 min timepoint. **(F)** GO-terms for upregulated genes in 60 min timepoint. **(G)** GO-terms for upregulated genes with CPEB-binding motifs in 10 min timepoint. **(H)** GO-terms for upregulated genes with CPEB-binding motifs in 60 min timepoint.

**Supplementary Table 4. High-confidence PASs. (A)** PASs predicted for datasets obtained 10 min after LTP induction by TAPAS. **(B)** High confidence poly(A) clusters predicted by LAPA for datasets obtained 10 min after LTP induction. **(C)** PASs predicted for datasets obtained 60 min after LTP induction. **(D)** High confidence poly(A) clusters predicted by LAPA for datasets obtained 60 min after LTP induction.

**Supplementary Table 5. Nonadenosine profiling upon LTP induction. (A)** Summary of Ninetails pipeline for dentate gyrus. **(B)** List of genes containing semi-templated poly(A) tails with their adjacent nucleotide contexts.

**Supplementary Table 6. Summary of synaptoneurosomal DRS/cDNA data per gene. (A)** Differential expression and differential adenylation data for unfractionated synaptoneurosomes DRS sequencing. (B) Summary of Ninetails pipeline for unfractionated synaptoneurosomes DRS sequencing. (C) Differential expression and differential adenylation data for monoribosome-bound mRNA synaptoneurosomes cDNA sequencing. (D) Differential expression and differential adenylation data for polyribosome-bound mRNA synaptoneurosomes cDNA sequencing. (E) Differential expression and differential adenylation data for unfractionated synaptoneurosomes cDNA sequencing.

## Supplementary Information

PDF file with Supplementary Figures.

### Description of additional data (provided via Zenodo)

Data accessible via link [https://doi.org/10.5281/zenodo.11478467] provided in Data availability section.

**CPEB1_motif.meme** - CPE1 motif sequence represented as position-dependent letter- probability matrice required by FIMO to make predictions.

**CPEB2_4_motif.meme** - CPE2,4 motif sequence represented as position-dependent letter-probability matrice required by FIMO to make predictions.

**mRatBN7.2_TAPAS_ref_flat.txt** - mRatBN7.2 reference annotation in format required by TAPAS

**mRatBN7.2_LAPA.gtf** - mRatBN7.2 reference annotation in format required by LAPA

**FIMO_output.zip** - compressed folder with motif predictions provided by FIMO software.

**LAPA_output_dentate_gyrus.zip** - compressed folder with poly(A) clusters predicted by LAPA software. Each timepoint is represented by separate output.

**TAPAS_output_dentate_gyrus.zip** - compressed folder with raw outputs produced by TAPAS software. Each timepoint is represented by separate output.

**Ninetails_output_dentate_gyrus.zip** - compressed folder with raw outputs produced by Ninetails software for samples from dentate gyrus. Subfolders are named according to the sample identifiers provided in Supplementary Table 1. For each sequencing run, 2 tsv files are produced: read classification and nonadenosine residue classification.

**Ninetails_output_synaptoneurosomes.zip** - compressed folder with raw outputs produced by Ninetails software for samples from synaptoneurosomes. Subfolders are named according to the sample identifiers provided in Supplementary Table 1. For each sequencing run, 2 tsv files are produced: read classification and nonadenosine residue classification.

**PolyA_clusters_high_confident.bed** - High-confidence poly(A) clusters annotation in bed format.

