## Supplementary Information for "Polyadenylation landscape of *in vivo* long-term potentiation in the rat brain"

For the manuscript entitled

#### Graphical abstract

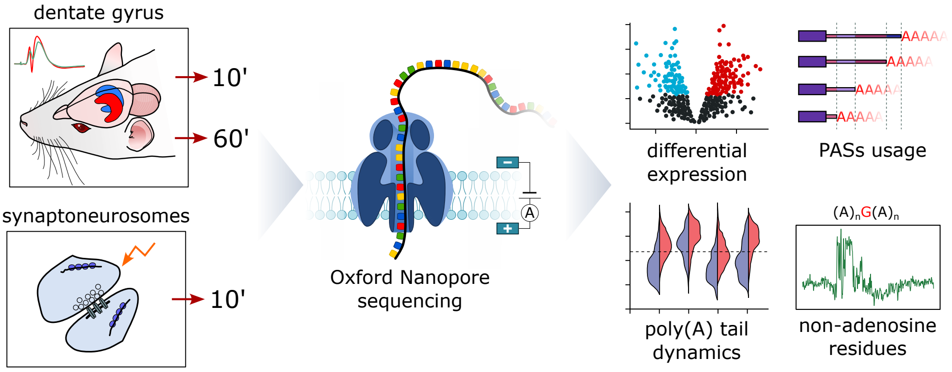

#### Supplementary Figure 1

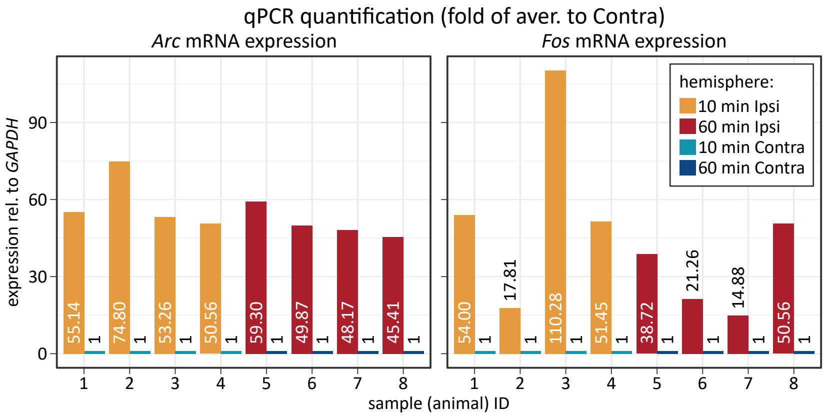

**Supplementary Figure 1. qPCR validation of LTP induction.**

Reverse transcription-quantitative PCR (RT-qPCR) analysis of *Arc* and *Fos* expression.

### Supplementary Figure 2

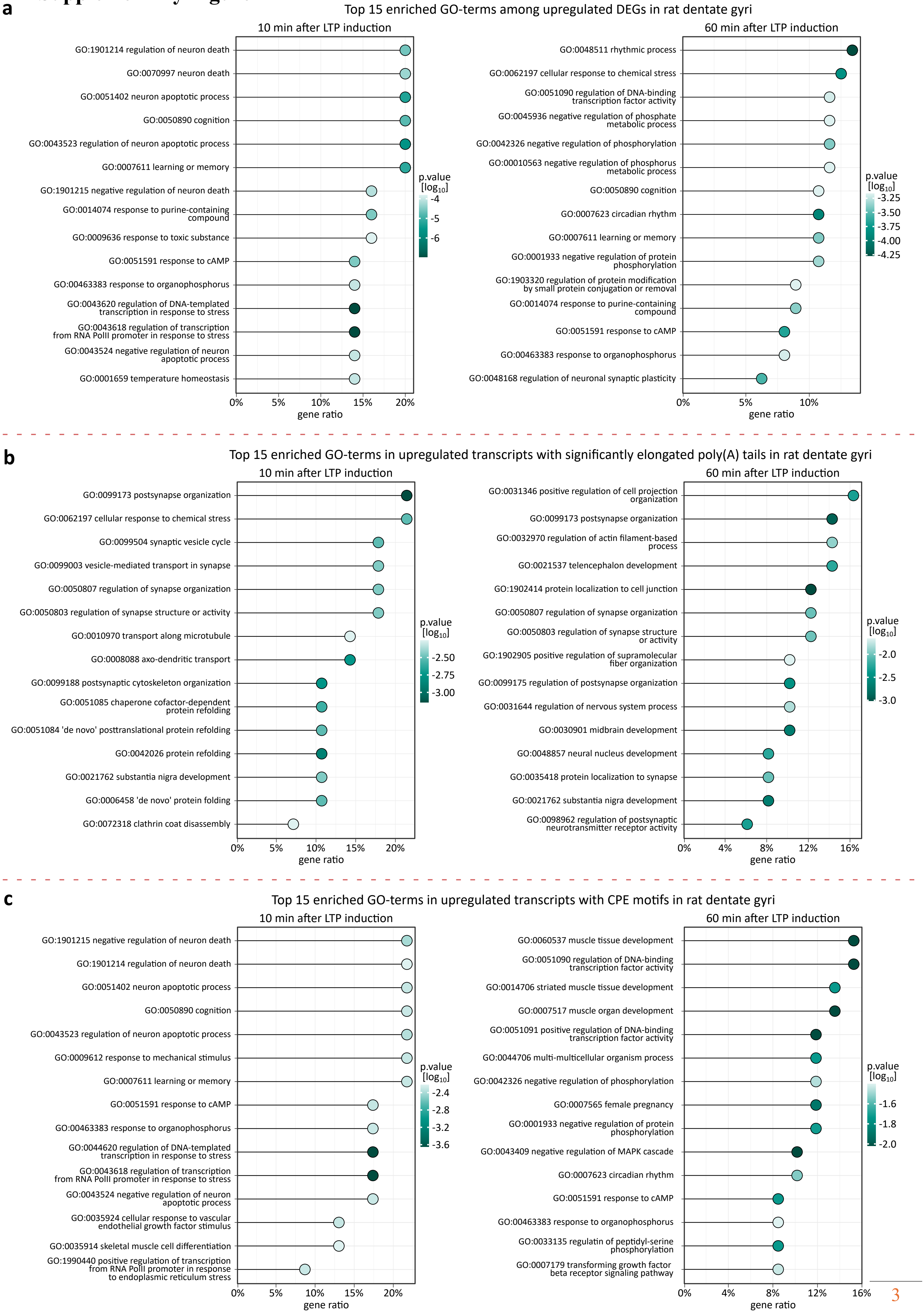

**Supplementary Figure 2.**

**Top 15 GO-terms of selected groups of transcripts ordered by gene ratio.**

GO-terms in dentate gyri shown for:

**a**, upregulated DEGs,

**b**, transcripts with significantly elongated poly(A) tails,

**c**, upregulated transcripts with CPE motifs.

### Supplementary Figure 3

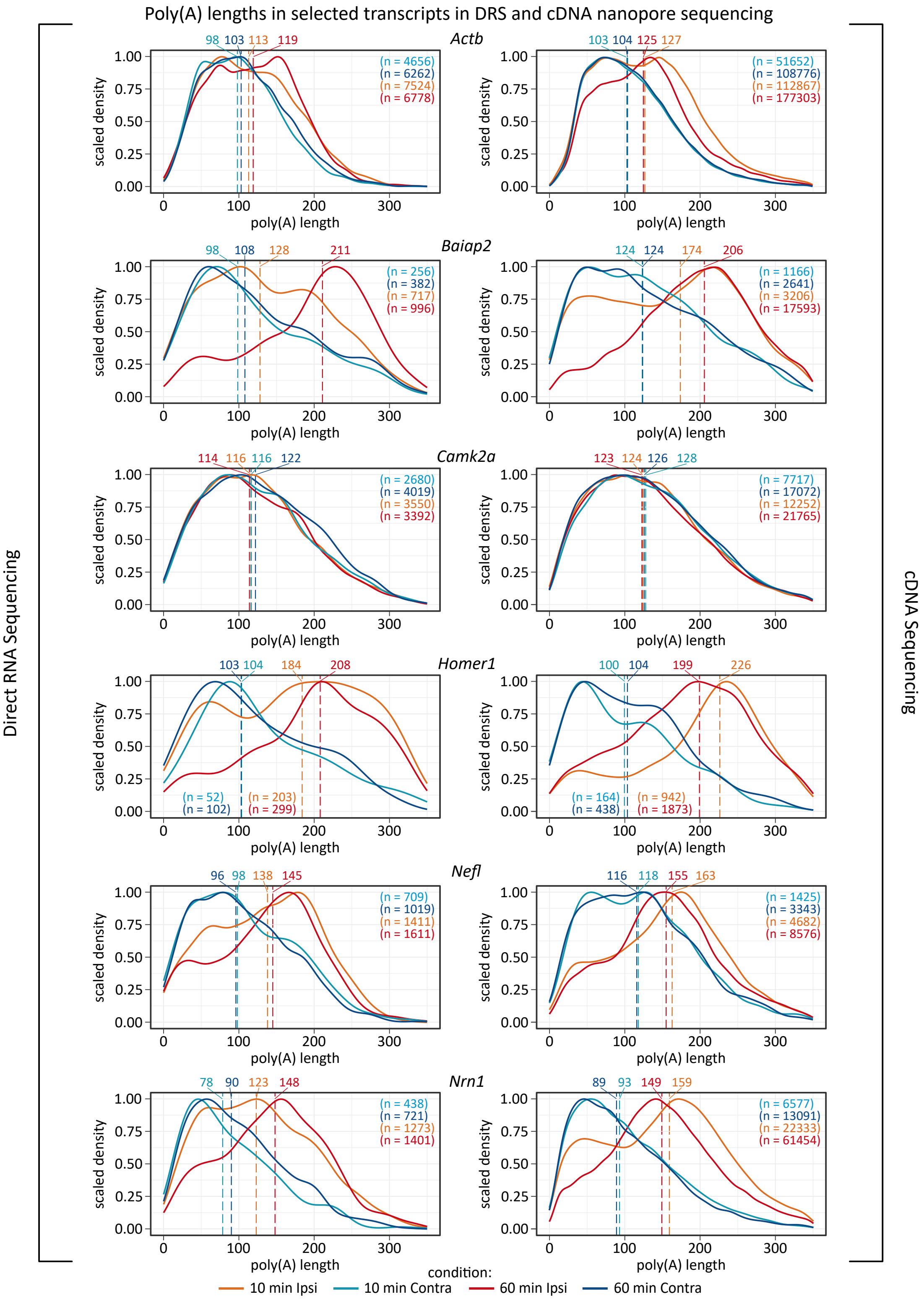

**Supplementary Figure 3.**

**Two nanopore sequencing methods used in this study provide consistent results.**

A comparison of poly(A) tail lengths distribution in selected transcripts reveals high concordance between Oxford Nanopore direct RNA sequencing (left) and cDNA sequencing (right) protocols.

Dashed lines indicate median tail length for each distribution; estimated median values in nucleotides are provided above the plotting area. Number of reads supporting each distribution are provided in brackets.

### Supplementary Figure 4

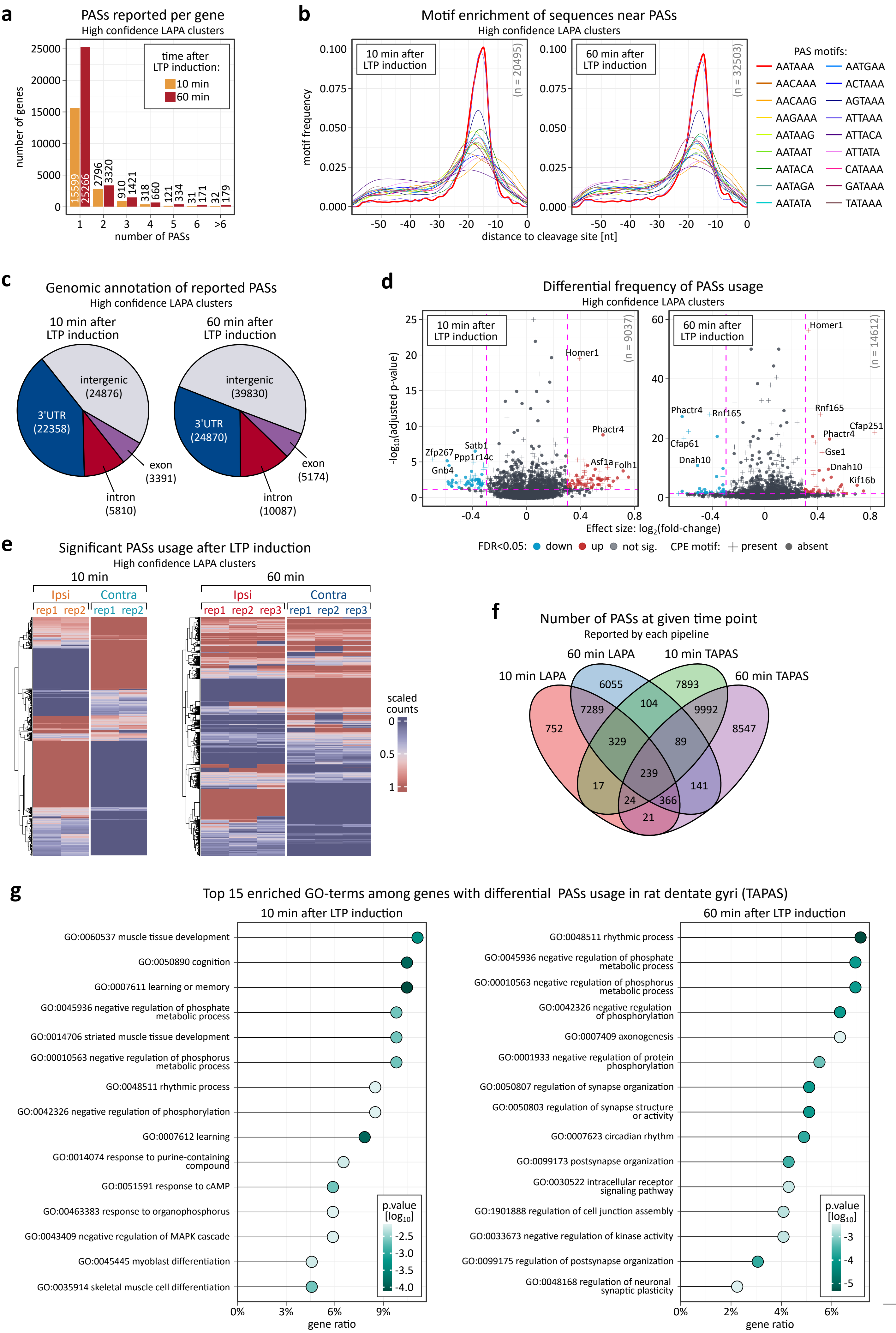

**Supplementary Figure 4.**

**Differential polyadenylation site usage in hippocampus upon LTP induction.**

**a**, Distribution of PASs reported per gene at 10 min and 60 min after LTP induction provided by LAPA.

**b**, Motif enrichment analysis of sequences near PASs at both time points, showing the frequency of known PAS motifs as reported by LAPA.

**c**, Genomic annotation of PASs detected at 10 min and 60 min after LTP induction, highlighting proportions in 3'UTR, exonic, intronic, and intergenic regions.

**d**, Volcano plots depicting differential PAS usage at 10 min and 60 min, with significant upregulated and downregulated PASs detected by LAPA. PAS is considered as differentially expressed if Δ usage is > 0.3. Differential usage of poly(A) sites was calculated using Fisher's exact test (two-sided, alpha = 0.05).

**e**, Heatmaps illustrating significant PAS usage changes in ipsilateral (Ipsi) and contralateral (Contra) hemispheres across replicates at both time points as provided by LAPA.

**f**, Venn diagram comparing PASs reported by LAPA and TAPAS pipelines at 10 min and 60 min after LTP induction.

**g**, Top 15 enriched GO-terms associated with genes exhibiting differential PASs usage, categorized by time point as reported by TAPAS.

### Supplementary Figure 5

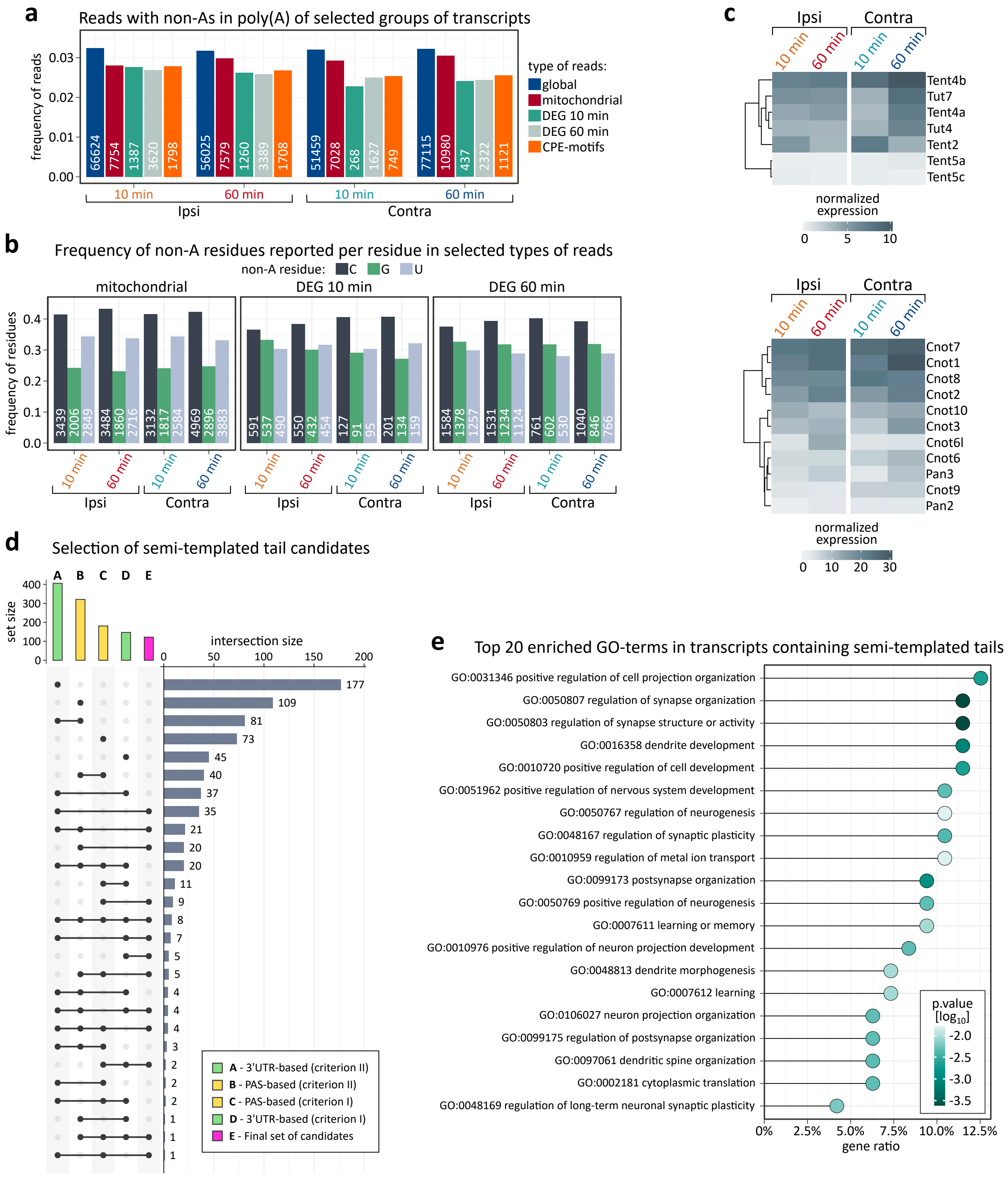

**Supplementary Figure 5.**

**Profiling of non-adenosine composition in hippocampus upon LTP induction.**

**a**, Frequency of reads with poly(A) tails decorated with non-adenosines. Data normalized on y-axis (each group separately).

**b**, Frequency of non-adenosine residues (cytidine, guanosine, and uridine, respectively) in selected groups of reads: mitochondrially-encoded genes and DEGs (each time point separately).

**c**, Expression of TENTs (upper heatmap) and deadenylases (bottom heatmap) in dentate gyrus.

**d**, Transcriptome-wide screening for semi-templated poly(A) tail candidates. Overlap between criteria used to identify transcripts with semi-templated tails and the intersection size of selected subsets.

**e**, Top 20 GO-terms of transcripts with semi-templated poly(A) tails ordered by gene ratio.

### Supplementary Figure 6

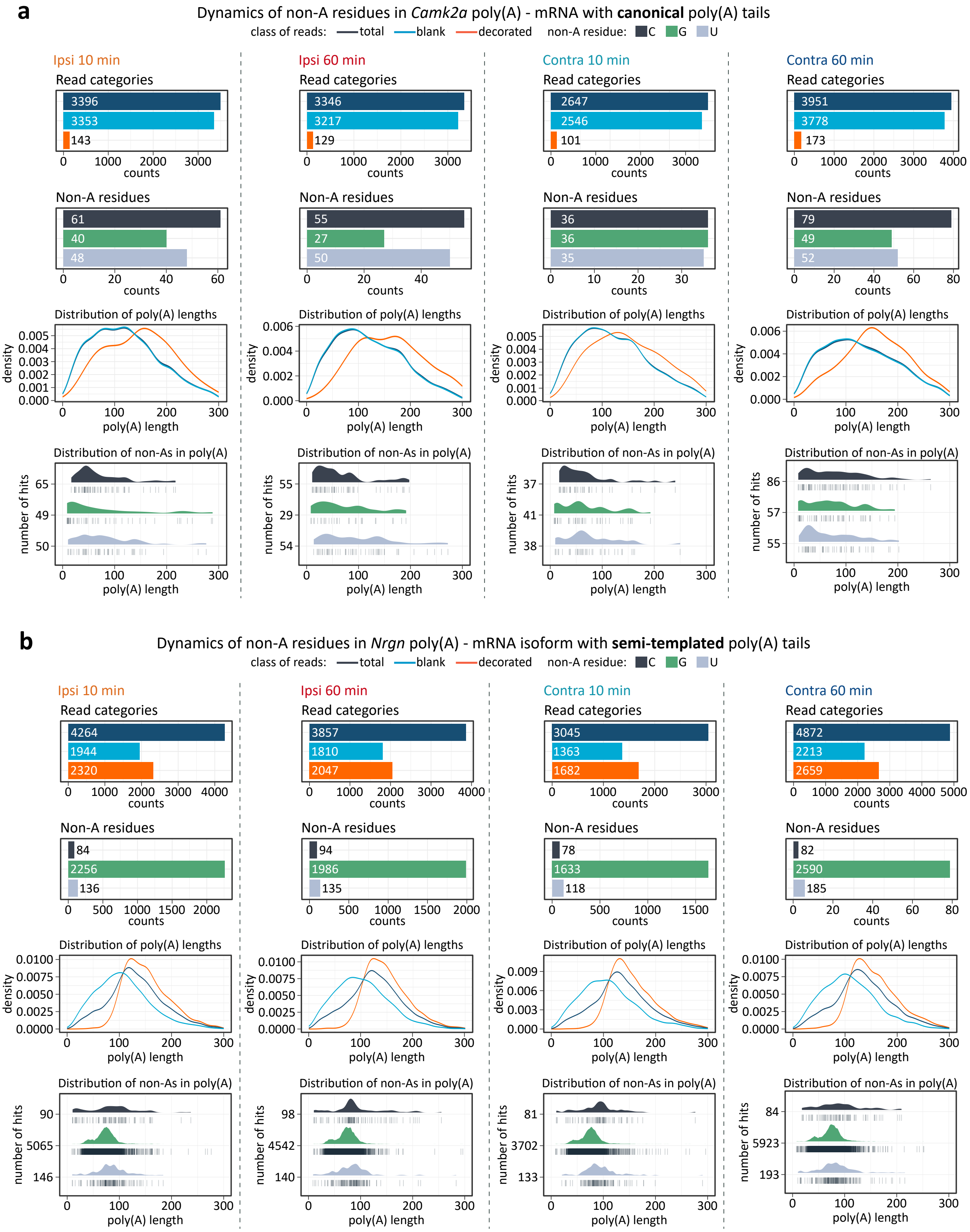

**Supplementary Figure 6.**

**Dynamics of non-adenosine residues in canonical vs. semi-templated poly(A) tails.**

**a**, Detailed overview of non-adenosine content in *Camk2a* poly(A) – an example of mRNA with canonical poly(A) tails across studied conditions (hemispheres, time points).

First row: frequency of read categories.
Second row: frequency of tails with given non-adenosines.
Third row: poly(A) tail length distribution of reads from either category.
Fourth row: estimated positions of non-adenosines within poly(A) tails.

In first, second and fourth rows, corresponding read counts (n) are provided.

**b**, Detailed overview of non-adenosine content in *Nrgn* poly(A) – an example of mRNA isoform with semi-templated poly(A) tails across studied conditions (hemispheres, time points).

First row: frequency of read categories.
Second row: frequency of tails with given non-adenosines.
Third row: poly(A) tail length distribution of reads from either category.
Fourth row: estimated positions of non-adenosines within poly(A) tails.

In first, second and fourth rows, corresponding read counts (n) are provided.

### Supplementary Figure 7

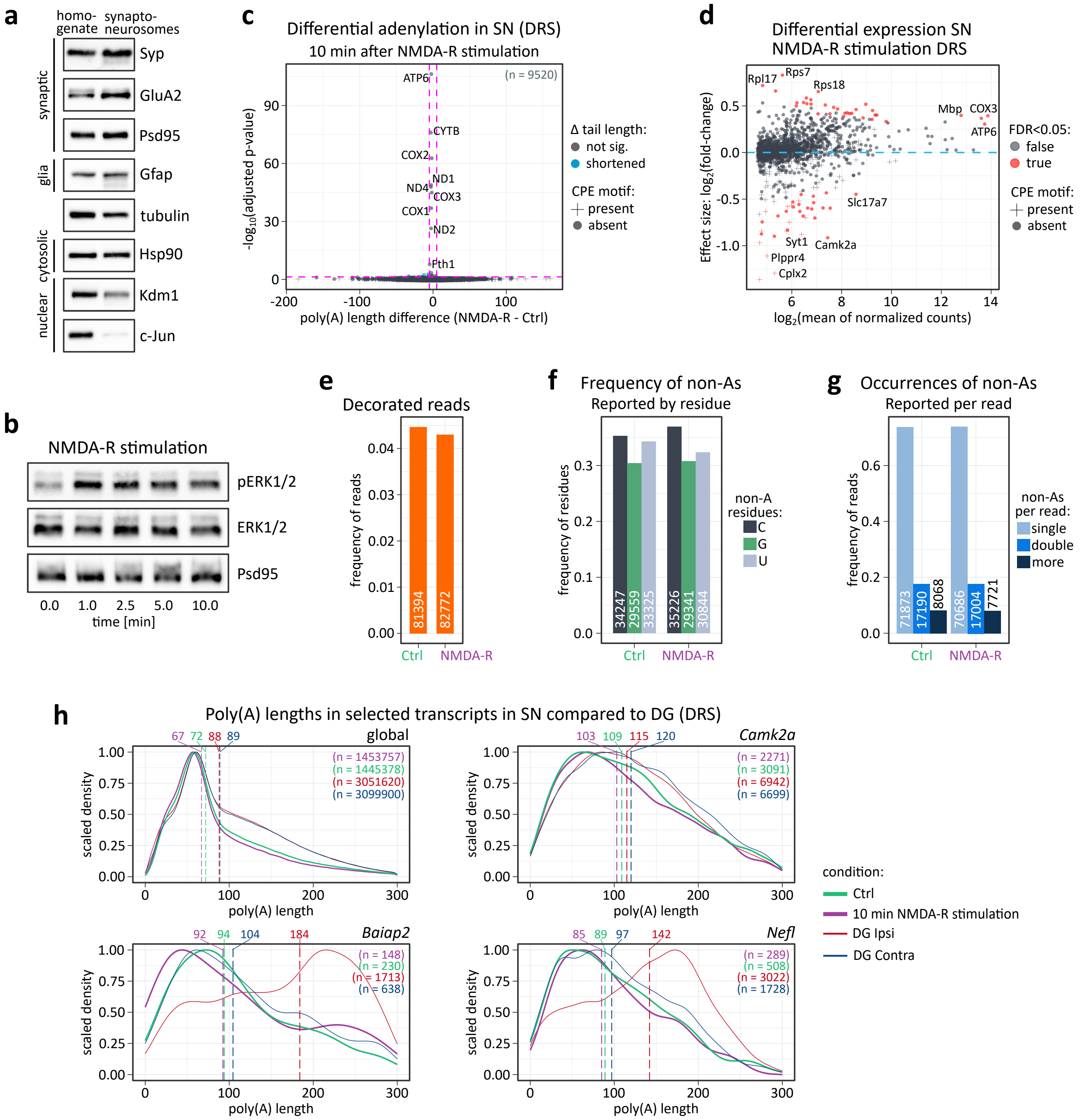

**Supplementary Figure 7.**

**Confirmation of synaptosome enrichment and investigation of nucleotide composition and length of synaptosomal poly(A) tails.**

**a**, Western blot validation of synaptoneurosome fractionation revealing the enrichment of both pre- and postsynaptic markers, but not glia (Gfap) and nuclear (Kdm1 and c-Jun) markers in the synaptoneurosomal fraction.

**b**, Western blot on synaptoneurosomes confirming activation of extracellular signal-regulated protein kinases 1 and 2 (ERK1/2) in response to the stimulation (0-10 min).

**c**, Polyadenylation profile of mRNAs in synaptoneurosomes upon NMDA-R stimulation. P.values between conditions were calculated using Wilcoxon signed-rank test, two-sided, alpha = 0.05, with Benjamini-Hochberg adjustment. The dashed lines divide the plot area into sectors based on p-value cut-off points and differences in poly(A) tail lengths. Unscaled version of Figure 6b.

**d**, DEGs in the synaptoneurosomes after NMDA-R stimulation. Differential expression was calculated with DESeq2. Data were shrunken (apeglm) for visualisation clarity.

**e**, Frequency of reads decorated with non-adenosines.

**f**, Occurrences of given non-adenosine (C, G, U) per read.

**g**, Reads decorated with given amount of non-adenosine instances.

**h**, Poly(A) tail length distribution in selected groups of transcripts in synaptoneurosomes vs dentate gyrus. The data from the given hemisphere in dentate gyrus were aggregated across time points for clarity. Dashed lines represent median tail lengths for each condition, with median values displayed above the plotting area. The total number of reads per condition (n) is also provided.

### Supplementary Figure 8

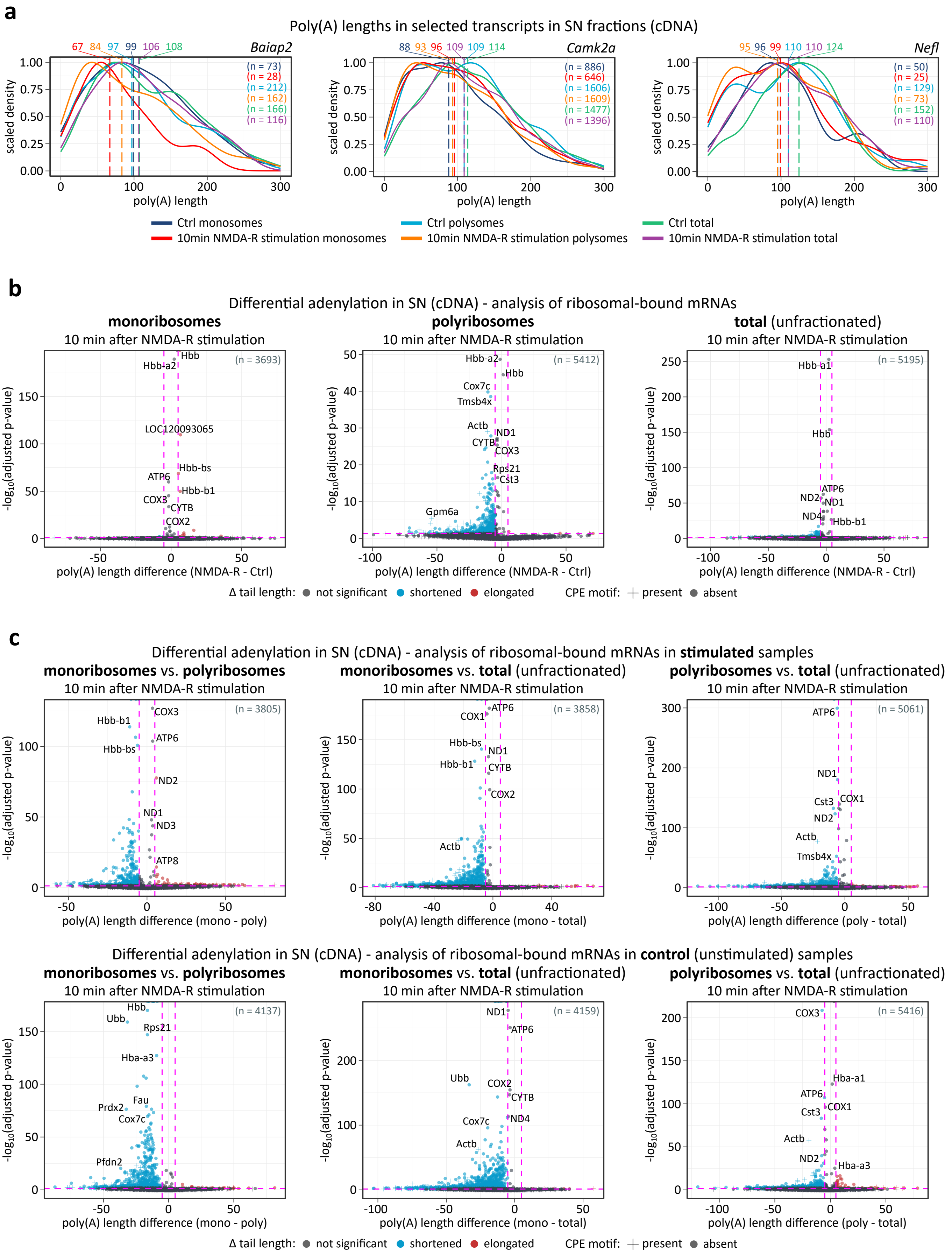

**Supplementary Figure 8.**

**The analysis of poly(A) tails of ribosomal-bound mRNA in synaptoneurosomes.**

**a**, Poly(A) tail length distribution in selected groups of transcripts in ribosomal-bound mRNA from synaptoneurosomes, cDNA, respectively: *Baiap2*, *Camk2a*, and *Nefl* – genes related to synaptic plasticity. Dashed lines represent median tail lengths for each condition, with median values displayed above the plotting area. The total number of reads per condition (n) is also provided.

**b**, Polyadenylation profile of ribosomal-bound mRNAs in synaptoneurosomes fractions upon NMDA-R stimulation. P.values between conditions were calculated using Wilcoxon signed-rank test, two-sided, alpha = 0.05, with Benjamini-Hochberg adjustment. Dashed lines divide the plot area into sectors based on p-value cut-off points and differences in poly(A) tail lengths. Unscaled plots (scaled version provided in Figure 6d).

**c**, Polyadenylation profile of ribosomal-bound mRNAs in synaptoneurosomes fractions upon NMDA-R stimulation – pairwise comparison between fractions. P.values between conditions were calculated using Wilcoxon signed-rank test, two-sided, alpha = 0.05, with Benjamini-Hochberg adjustment. Dashed lines divide the plot area into sectors based on p-value cut-off points and differences in poly(A) tail lengths. Unscaled plots. Upper row – stimulated samples, bottom row – unstimulated controls.
